# An integrative skeletal and paleogenomic analysis of prehistoric stature variation suggests relatively reduced health for early European farmers

**DOI:** 10.1101/2021.03.31.437881

**Authors:** Stephanie Marciniak, Christina M. Bergey, Ana Maria Silva, Agata Hałuszko, Mirosław Furmanek, Barbara Veselka, Petr Velemínský, Giuseppe Vercellotti, Joachim Wahl, Gunita Zariņa, Cristina Longhi, Jan Kolář, Rafael Garrido-Pena, Raúl Flores-Fernández, Ana M. Herrero-Corral, Angela Simalcsik, Werner Müller, Alison Sheridan, Žydrūnė Miliauskienė, Rimantas Jankauskas, Vyacheslav Moiseyev, Kitti Köhler, Ágnes Király, Beatriz Gamarra, Olivia Cheronet, Vajk Szeverényi, Viktoria Kiss, Tamás Szeniczey, Krisztián Kiss, Zsuzsanna K. Zoffmann, Judit Koós, Magdolna Hellebrandt, László Domboróczki, Cristian Virag, Mario Novak, David Reich, Tamás Hajdu, Noreen von Cramon-Taubadel, Ron Pinhasi, George H. Perry

## Abstract

Human culture, biology, and health were shaped dramatically by the onset of agriculture ~12,000 years before present (BP). Subsistence shifts from hunting and gathering to agriculture are hypothesized to have resulted in increased individual fitness and population growth as evidenced by archaeological and population genomic data alongside a simultaneous decline in physiological health as inferred from paleopathological analyses and stature reconstructions of skeletal remains. A key component of the health decline inference is that relatively shorter statures observed for early farmers may (at least partly) reflect higher childhood disease burdens and poorer nutrition. However, while such stresses can indeed result in growth stunting, height is also highly heritable, and substantial inter-individual variation in the height genetic component within a population is typical. Moreover, extensive migration and gene flow were characteristics of multiple agricultural transitions worldwide. Here, we consider both osteological and ancient DNA data from the same prehistoric individuals to comprehensively study the trajectory of human stature variation as a proxy for health across a transition to agriculture. Specifically, we compared ‘predicted’ genetic contributions to height from paleogenomic data and ‘achieved’ adult osteological height estimated from long bone measurements on a per-individual basis for n=160 ancient Europeans from sites spanning the Upper Paleolithic to the Iron Age (~38,000-2,400 BP). We found that individuals from the Neolithic were shorter than expected (given their individual polygenic height scores) by an average of −4.47 cm relative to individuals from the Upper Paleolithic and Mesolithic (P=0.016). The average osteological vs. expected stature then increased relative to the Neolithic over the Copper (+2.67 cm, P=0.052), Bronze (+3.33 cm, P=0.032), and Iron Ages (+3.95 cm, P=0.094). These results were partly attenuated when we accounted for genome-wide genetic ancestry variation in our sample (which we note is partly duplicative with the individual polygenic score information). For example, in this secondary analysis Neolithic individuals were −3.48 cm shorter than expected on average relative to individuals from the Upper Paleolithic and Mesolithic (P=0.056). We also incorporated observations of paleopathological indicators of non-specific stress that can persist from childhood to adulthood in skeletal remains (linear enamel hypoplasia, cribra orbitalia, and porotic hyperostosis) into our model. Overall, our work highlights the potential of integrating disparate datasets to explore proxies of health in prehistory.

## Introduction

The agricultural revolution – beginning ~12,000 BP (years before present) in Mesopotamia^1,2^ and then spreading^3–5^ or occurring independently^6,7^ across much of the inhabited planet – precipitated profound changes to human subsistence, social systems, and health. Seemingly paradoxically, the agricultural transition may have presented conflicting biological benefits and costs for early farming communities^8,9^. Specifically, demographic reconstructions from archaeological and population genetic records suggest that the agricultural transition led to increased individual fitness and population growth^6,10–12^, likely due in part to new food production and storage capabilities. Yet, bioarchaeological analyses of human skeletal remains from this cultural period suggest simultaneous declines in individual physiological well-being and health, putatively from i) nutritional deficiency and/or ii) increased pathogen loads as a function of greater human population densities, sedentary lifestyles, and proximity to livestock^9,13–18^.

To date, anthropologists have used two principal approaches to study health across the foraging-to-farming transition in diverse global regions^13,19,20^. The first approach involves identifying paleopathological indicators of childhood stress that persist into adult skeletal remains. For example, porotic hyperostosis (porous lesions on the cranial vault) and cribra orbitalia (porosity on the orbital roof) reflect a history of bone marrow hypertrophy or hyperplasia resulting from one or more periods of infection, metabolic deficiencies, malnutrition, and/or chronic disease^21–26^. Meanwhile, linear enamel hypoplasia (transverse areas of reduced enamel thickness on teeth) occurs in response to similar childhood physiological stressors (e.g., disease, metabolic deficiencies, malnutrition, weaning) that disrupt enamel formation in the developing permanent dentition^27–30^. Broadly, these paleopathological indicators of childhood stress tend to be observed at higher rates among individuals from initial farming communities relative to earlier periods, potentially reflecting their overall “poorer” health^14,31–36^.

A second approach uses skeleton-based estimates of achieved adult stature as a proxy for health during childhood growth and development^37–39^. Since stature is responsive to the influences of nutrition and disease burden alongside other factors, relatively short “height-for-age” (or “stunting”) has been used as an indicator of poorer health in both living and bioarchaeological contexts^39–43^. When studying the past, individual stature can be estimated from long bone measurements and regression equations^44–47^. Using these methods, multiple prior studies have reported a general profile of relatively reduced stature for individuals from early agricultural societies in Europe^15,36,48–50^, North America^51–53^, the Levant^16,32^, and Asia^54,55^. For example, estimated average adult mean statures for early farmers are ~10 cm shorter relative to those for preceding hunter-gatherers in both Western Europe (females −8 cm; males −14 cm)^49,50^ and the Eastern Mediterranean (females −11 cm; males −8 cm)^56^. This pattern is not universal, as a few studies do not report such changes^57,58^; the variation could be informative with respect to identifying potential underlying factors^59^.

However, in addition to environmental effects like childhood nutrition and disease, inherited genetic variation can have an outsized impact on terminal stature, with ~80% of the considerable degree of height variation within many modern populations explainable by heritable genetic variation^60–63^. Moreover, migration and gene flow likely accompanied many subsistence shifts in human prehistory. For example, there is now substantial paleogenomic evidence of extensive population turnover across prehistoric Europe^64–69^. Therefore, from osteological studies alone we are unable to quantify the extent to which temporal changes in height reflect variation in childhood health versus changes/differences in the frequencies of alleles associated with height variation.

In this study we have performed a combined analysis of ancient human paleogenomic and osteological data where both are available from the same n=160 prehistoric European individuals representing cultural periods from the Upper Paleolithic (~38,000 BP) to the Iron Age (~2,400 BP). This approach allows us to explore whether ‘health’, as inferred from the per-individual difference between predicted genetic contributions to height and osteological estimates of achieved adult height, changed over the Neolithic cultural shift to agriculture in Europe. When craniodental elements were preserved and available for analysis (n=91 of the 160 individuals), we also collected porotic hyperostosis, cribra orbitalia, and linear enamel hypoplasia paleopathological data in order to examine whether patterns of variation between osteological height and genetic contributions to height are explained in part by the presence/absence of these indicators of childhood or childhood-inclusive stress.

## Results

We developed a database of n=160 ancient European adult individuals (65 females, 95 males) with available genome-wide paleogenomic data (either from shotgun sequencing or DNA capture-based approaches; and from both published and in-process studies) and stature estimates based on long bone measurements (either newly collected or published; Fig. 1A; *SI Appendix*, Table S1). The cultural and time periods represented in our dataset are: Upper Paleolithic (38,000-12,000 BP; n=8 individuals), Mesolithic (11,000-6,400 BP; n=16), Neolithic (7,100-3,500 BP; n=39), Copper Age (6,300-3,400 BP; n=58), Bronze Age (4,500-2,500 BP; n=32), and Iron Age (2,600-2,400 BP; n=7).

**Figure 1.**
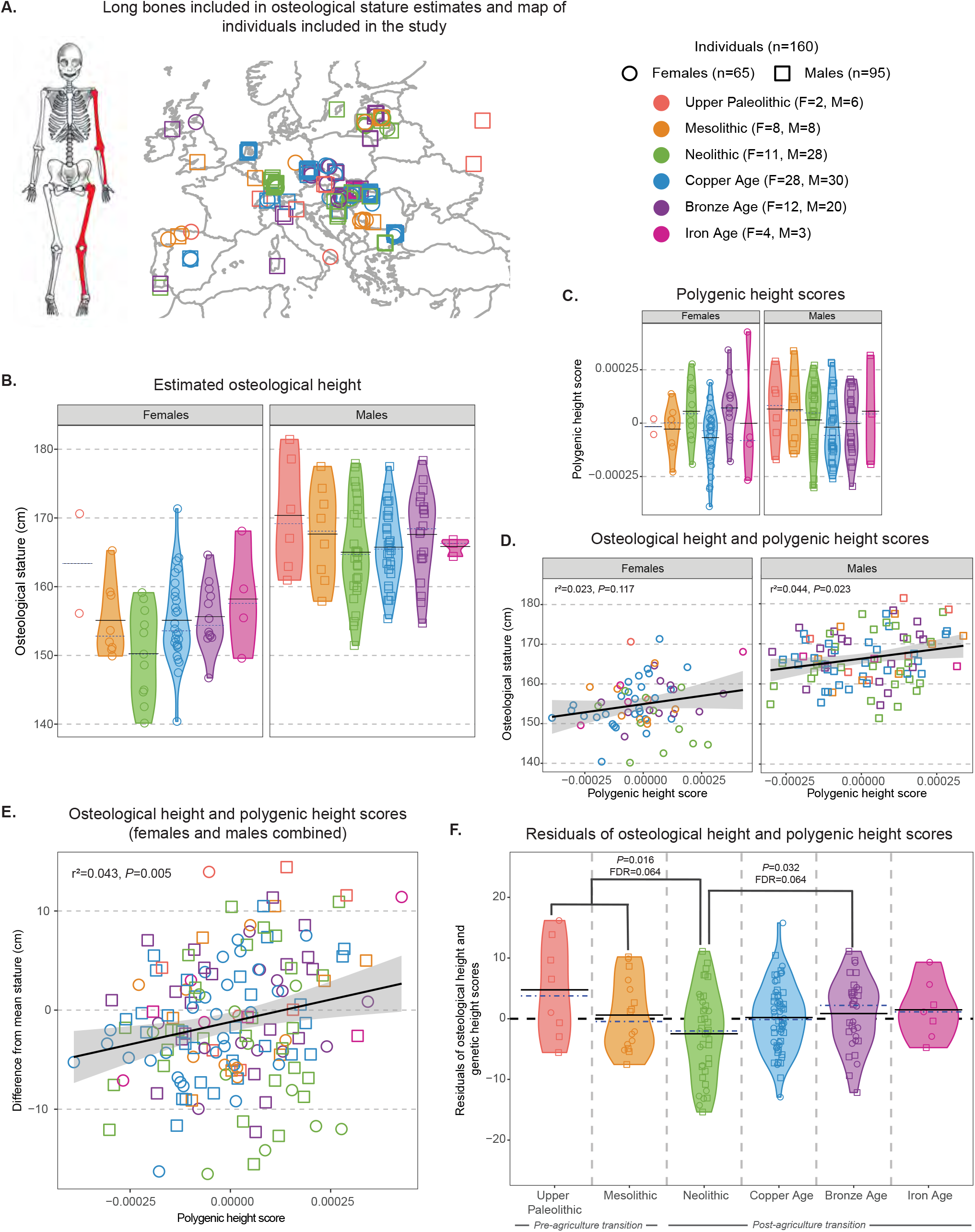
Osteological stature and ancient DNA-based polygenic height scores. **A)** Map of the locations of the archaeological sites from which individuals included in data set were recovered. **B)** Osteological height estimates generated using measurements of long bone lengths (highlighted in red on the illustration) and sex-specific regression equations (Ruff et al., 2012)^44^. **C)** Polygenic height scores generated using genome-wide association summary statistics for height-associated single nucleotide polymorphisms and individual ancient DNA genotype data. **D,E)** The relationship between polygenic height score and estimated osteological stature (cm) for females, males, and for the full sample with height differences from mean stature calculated separately for females (mean=156.45 ± 4.32 cm) and males (mean=167.04 ± 1.95 cm), respectively. **F)** Residuals of the relationship between polygenic height score and osteological height with sex as a covariate for all individuals, by cultural period. Mean and median are represented by the black and blue dashed lines, respectively. Skeletal illustration by Katharine Thompson.

Subsistence strategies in the Upper Paleolithic and Mesolithic focused on gathering, collecting, and hunting food. The Neolithic is marked by the emergence of plant cultivation and animal domestication (to varying degrees and tempos of integration), long-term settlements, larger populations, and increased social complexity – processes which then intensified and expanded in subsequent periods^70^. The overlapping dates among the different cultural periods reflect both geographical variation in the timing of cultural change and the potential for co-occurrence of multiple cultural traditions within a single region.

### Confirmation of an average osteological stature dip in the Neolithic

We used an osteometric board to newly estimate the lengths of preserved long bones for n=86 of the n=160 total individuals (53.7%) in our database *(SI Appendix*, Table S1). We also recorded published and unpublished (previously collected) long bone length estimates for an additional n=54 individuals (33.7%). In these cases we estimated osteological stature from the long bone length data^44^. Finally, for n=20 individuals only pre-calculated terminal height estimates were available (12.5%).

We observed osteological stature variation among cultural periods (Figure 1B). Reconstructions of osteological stature for females and males are lower during the initial shift to farming during the Neolithic compared with earlier and post-Neolithic periods. Specifically, individuals from the pre-Neolithic periods (female average stature = 156.7 cm ± 6.8 (s.d.) cm, males=168.8 ± 7.4 cm) were ~4 cm taller on average than those from the Neolithic (females=150.3 ± 6.6 cm; males 165.0 ± 7.2 cm; linear model including sex as a factor; P=0.017; FDR=0.109). Then, relative to the Neolithic, average osteological stature rebounded in our Copper Age (females=155.1 ± 6.3 cm; males=165.7 ± 5.4 cm; P=0.131; FDR=0.257), Bronze Age (females=155.6 ± 4.8 cm; males 167.6 ± 6.3 cm; P=0.031; FDR=0.109), and Iron Age (females=158.2 ± 7.8 cm; males=165.8 ± 1.3 cm; P=0.147; FDR=0.257) samples *(SI Appendix*, Table S2a, and S2b).

The overall pattern from our data roughly parallels previously-published reports. Specifically: a) stature decreased slightly from the Upper Paleolithic to the Mesolithic^35,36,71^; b) marked stature reduction occurred during the initial agricultural transition in the Neolithic^14,16,34,55^ (although this is not universal^70,72–74^); and c) stature rebounded during subsequent post-Neolithic periods of agricultural intensification^14,16^.

### Early farmers were relatively shorter than expected given their polygenic height scores

We next considered the osteological height estimates in the context of ancient DNA-based polygenic height scores for the same individuals. Using an established approach for working with ancient DNA genotype data^65,75,76^, we estimated a polygenic score for each prehistoric individual based on their available genome-wide genotypes in the context of results from a large-scale genome-wide association study of stature variation in modern Europeans^77^ (data from: http://www.nealelab.is/uk-biobank/). For the results presented in the main text and figures, we used a version of the dataset in which all variants possibly affected by deamination-based ancient DNA damage^78–80^ were masked. We also performed all analyses with the unmasked dataset and obtained similar results (*SI Appendix*, Fig. S1, Table S3).

While the polygenic scores that we estimated for the n=160 individuals were somewhat variable across cultural periods (Figure 1C; *SI Appendix*, Table S4 and S5) as expected based on prior work^58^, we were most interested in using these data to begin to account for genetic contributions to achieved adult (osteological) height on a per-individual basis. Polygenic height scores and osteological estimates of stature were positively correlated for females (n=65; Fig. 1D; r^2^=0.023; P=0.117), males (n=95; Fig. 1D; r^2^=0.044; P=0.023), and for the combined dataset when calculating the difference between per-individual osteological height and mean stature separately for females (mean=156.45 ± 4.32 cm) and males (mean=167.04 ± 1.95 cm) (Fig. 1E; r^2^=0.043 P=0.005). These results support the general biological plausibility of our integrative analysis of paleogenomic and osteological data.

Importantly and expectedly, there is still considerable inter-individual variation in the relationship between polygenic height score and achieved adult stature, which could reflect any combination of incomplete genetic information, long bone measurement error, polygenic height score or stature estimate error, and the effects of childhood nutrition, disease, and other environmental variables on growth. Accordingly, we next analyzed the residuals from the combined-sex osteological stature and ancient DNA-based polygenic score model (Fig. 1E) to test whether individuals tended to have taller or shorter adult stature relative to expectations given their individual polygenic scores across the different cultural periods. These residuals are expressed in +/- cm from predicted stature per individual.

When using our ancient DNA-based approach to partly account for the predicted contribution of genetic variation to adult stature, we observed that individuals from the Neolithic were indeed osteologically shorter than expected (i.e. based on their polygenic height scores, and in the context of our overall sample) compared to individuals from other cultural periods (Fig. 1F; *SI Appendix*, Table S6). Specifically, pre-Neolithic individuals (average residual = +1.99 ± 6.8 cm) were +4.47 cm taller than expected on average relative to Neolithic individuals (average residual = −2.48 ± 9.9 cm; P=0.016; FDR=0.064). The average osteological vs. genetic height score residual then increased steadily in the Copper Age (+2.67 cm relative to the Neolithic; P=0.052; FDR=0.069), Bronze Age (+3.33 cm; P=0.032; FDR=0.064), and Iron Age (+3.95 cm; P=0.094; FDR=0.094).

We confirmed that these results cannot be explained by geographic variation (latitude and longitude) in our sample *(SI Appendix*, Fig. S2, Table S7). We also obtained similar results when we separately analyzed females and males (*SI Appendix*, Fig. S3, Table S6) and when we separately analyzed the lengths of individual long bones as opposed to the reconstructed stature estimates (for the femur and radius, in particular, although there are sample size limitations) (*SI Appendix*, Fig. S4 and S5, Table S8).

In contrast, our results were partially muted when we included variables explicitly reflecting genetic ancestry in our model. Our primary approach (as above) already accounts for ancestry variation via individual-level calculations of polygenic scores. However, polygenic height scores explain only a proportion of total heritable variation^81–83^. Therefore, we repeated our hypothesis tests after conducting a multi-dimensional scaling (MDS) analysis with the genome-wide genotype data for all n=160 ancient individuals to then including the first four MDS axes from this analysis as factors in an updated linear model, following Cox et al.^84^. After doing so, pre-Neolithic individuals (average residual = +2.31 ± 6.8 cm) were +3.48 cm taller than expected on average relative to Neolithic individuals (average residual = −1.16 ± 6.9 cm; P=0.056, FDR=0.222). The trend towards increasing average values through the Copper Age (+0.88 cm relative to the Neolithic; P=0.503; FDR=0.671), Bronze Age (+1.46 cm; P=0.310; FDR=0.620), and Iron Age (+0.62 cm; P=0.700; FDR=0.770) still exists but is considerably less pronounced (*SI Appendix*, Fig. S6, Table S9).

### Paleopathological indicators of non-specific stress

Adverse early life conditions may negatively impact adult stature. To begin to investigate whether individual-level early life effects on prehistoric stature could be identified, we incorporated observations of paleopathological indicators of non-specific stress that can persist from childhood to adult skeletal remains into our analytical model. To do so, we characterized the presence/absence of one or more of cribra orbitalia (porosity on the orbital roof), porotic hyperostosis (porosity on the cranial vault), and linear enamel hypoplasia (reduced areas of enamel thickness) for n=91 of the n=160 (56.9%) individuals in our study (n=75 newly characterized; n=16 published/previously characterized *SI Appendix*, Table S10).

For 53 of these 91 individuals (58.2%), crania were sufficiently complete for assessment of the presence/absence of all three stress indicators (Fig. 2A). Of this subsample, one or more stress indicators were present for 39 (73.6%) of the individuals, two or more indicators were observed in 18 (34%) individuals, and all three paleopathological indicators were present in only two (3.8%) individuals. Thus, stresses on health were relatively common overall in prehistoric Europe.

**Figure 2.**
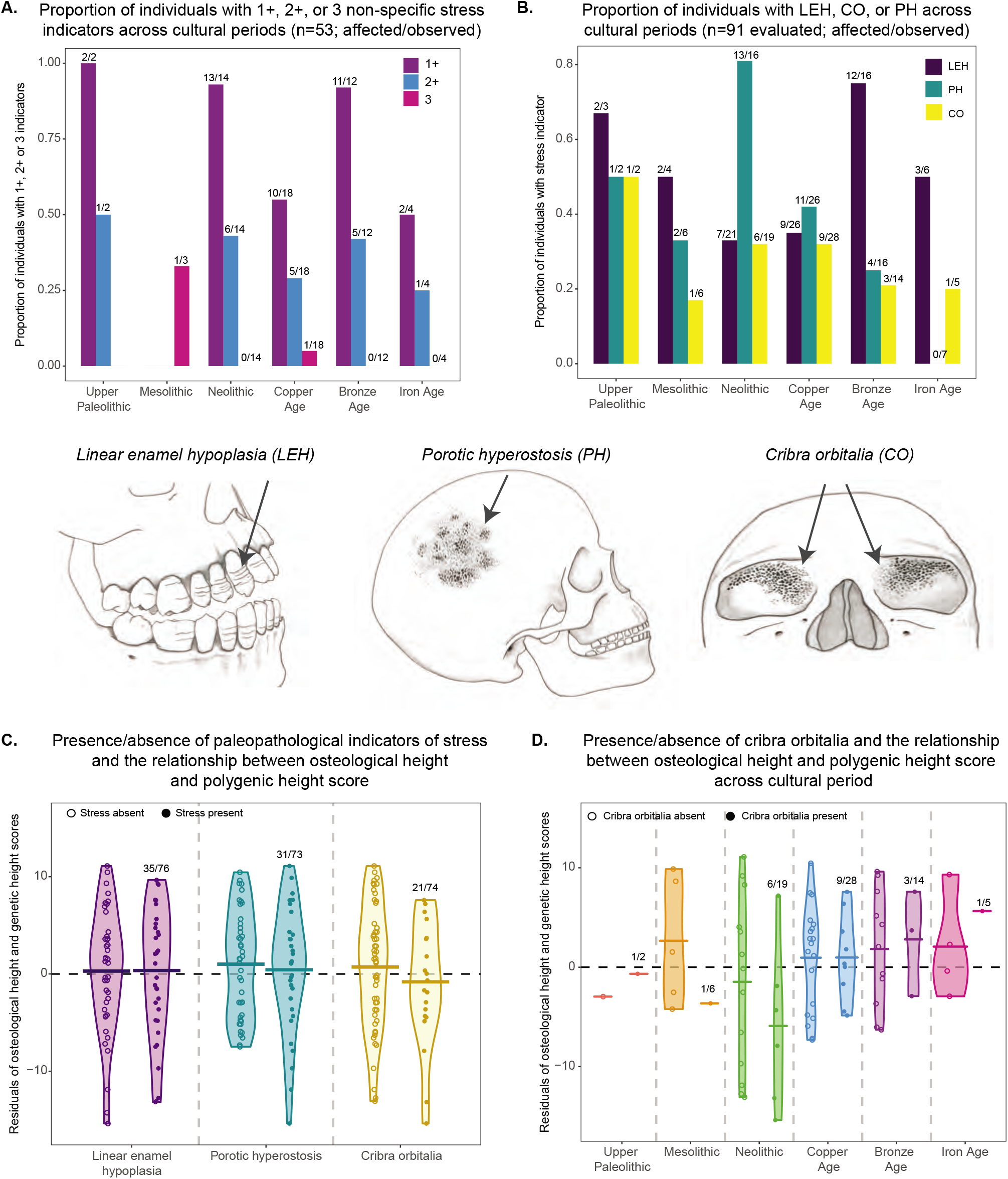
Paleopathological indicators of stress. Paleopathological indicators of non-specific stress evaluated in this study: linear enamel hyopolasia (LEH; bands of reduced enamel thickness on teeth), cribra orbitalia (CO; porosity on the orbits), and porotic hyperostosis (PO; porosity on the side of the skull). **A)** The remains of 53 individuals were sufficiently complete to permit who presence/absence assessment for all three paleopathologies. The proportions of individuals with 1+, 2+, and all 3 stress indicators are indicated, across cultural periods. Numbers above the bars indicate sample sizes. **B)** The presence/absence of at least one of the three paleopathological indicators of stress could be determined for 91 total individuals. Shown are the proportions of individuals (of those who could be assessed for that indicator) with LEH, CO, and PH, across cultural periods. Numbers above the bars indicate sample sizes. **C)** Residuals of the relationship between polygenetic height score and osteological height with sex as a covariate plotted separately for individuals with each paleopathological indicator of stress present vs. absent. Means are represented by the black lines. Numbers above the bars indicate sample sizes. **D)** Residuals of the relationship between polygenetic height score and osteological height with sex as a covariate plotted separately for individuals with and without cribra orbitalia, by cultural period. Means are represented by the black lines. Numbers above the bars indicate sample sizes. Skeletal illustrations by Katharine Thompson.

A striking 92.9% (13/14) of Neolithic individuals had one or more stress indicators (Fig. 2A). While the proportion of Copper Age individuals with one or more stress indicators (10/18; 55.6%) was lower compared to that for the Neolithic (Fisher’s exact test; P=0.044), the Neolithic result is not unique, with one or more stress indicators also recorded for all but one individual in the Bronze Age sample (11/12; 91.7%).

Considering the larger dataset of n=91 individuals with presence/absence data for at least one stress indicator to maximize sample sizes, we observed a distinct difference between the Neolithic and Bronze Age patterns (Fig. 2B). Specifically, porotic hyperostosis is common in the Neolithic sample (13/16; 81.3%) while linear enamel hypoplasia is relatively rare (7/21 individuals; 33.3%). The opposite is true in the Bronze Age, with 4/16 (25%) positive for porotic hyperostosis yet 12/16 (75%) with linear enamel hypoplasia.

We next tested whether the presence of paleopathological indicators of stress is predictive of individual-level deviations from the overall relationship between osteological stature and polygenic height score estimates. Based on the subset of individuals with presence/absence data for all three paleopathological indicators, the 39 individuals with one or more stress indicator were −1.36 cm shorter than expected on average compared to the 14 individuals without any stress indicators (*SI Appendix*, Fig. S7, Table S11). With the moderate magnitude of this difference and the relatively small sample size available for this analysis, this result was not unlikely based on chance expectations (t-test; P=0.448). The 18 individuals with two or more stress indicators were −1.80 cm shorter than expected on average relative to individuals with no stress indicators (P=0.411).

Using our larger dataset, we next separately analyzed the effect of each paleopathological stress indicator on the osteological stature-polygenic height score relationship (Fig. 2C). We found that the n=21 individuals with cribra orbitalia were slightly but not significantly shorter (−1.52 cm) than expected on average compared to the n=53 individuals without cribra orbitalia (P=0.361; FDR=0.968). Linear enamel hypoplasia and porotic hyperostosis presence/absence were negligibly associated with osteological stature vs. polygenetic height score residual variation (Fig. 2D; *SI Appendix*, Table S12a). These patterns did not change appreciably when excluding the few individuals with active cribra orbitalia or porotic hyperostosis lesions from their respective analyses (active lesions reflect adult stress while potentially masking evidence of lesions from childhood; *SI Appendix*, Fig. S8, Table S12b).

Finally, we investigated the relationship between cribra orbitalia, porotic hyperostosis, and linear enamel hypoplasia presence/absence and osteological vs. polygenic height score residual in the context of cultural period (*SI Appendix*, Fig. S9, Table S13, Table S14). Of note: the n=6 Neolithic individuals with cribra orbitalia were −4.44 cm shorter than expected on average compared to the n=13 individuals from the same cultural period without cribra orbitalia (Fig 2D; P=0.306). This preliminary but suggestive effect was nearly absent in the Copper Age (−0.015 cm difference; P=0.994), the only other period with sufficient presence/absence sample sizes for analysis.

## Discussion

Bioarchaeologists have equated repeated observations of relatively shorter average adult statures in the Neolithic to a likely general health decline for individuals during this cultural period^13,14,16,34,55,85^. Combinations of reduced nutritional diversity, unpredictable food availability (e.g., crop failure, storage loss), and increased infectious disease burden may have negatively impacted childhood health and growth^18,86,87^. Understandably, those prior studies did not account for the contribution of inter-individual variation in the contribution of heritable genetic factors to adult stature. Yet this consideration is especially important in light of updated understandings of considerable migration and gene flow processes associated with various farming transitions^88–91^.

In our study we sampled 160 prehistoric European individuals for whom both genome-wide ancient DNA data and intact long bones were available for analysis, making it possible for the first time to test whether Neolithic individuals were still osteologically shorter than expected when accounting (at least partly) for individual-level genetic contributions to height. Using this approach we found that the average Neolithic farmer was indeed relatively shorter than expected compared to pre-Neolithic individuals (Fig. 1F). Average osteological versus expected stature then increased over each post-Neolithic cultural period. This gradual recovery may reflect a history of continued (although variable) cultural and technological innovations that ameliorated and/or overpowered the initial nutritional and disease stressors faced by the earliest farmers^66–69,92,93^.

Our framework is related to but differs from that of a previous study by Cox and colleagues^58^, who compared population-level osteological height estimates (n=1,159 total individuals; the osteological data were from Niskanen et al.^94^ and ancient DNA-based polygenic height scores (n=1,071) across prehistoric Europe. These two estimates were computed separately, i.e. typically not for the same individuals, thereby facilitating the large sample sizes. In contrast, our approach expressly considers individual-level dynamics in the relationship between these two variables, which is sample size restrictive yet potentially insightful. Interestingly, mean osteological stature estimates and polygenic height scores were both similar between the European Mesolithic and Neolithic^58,94^. This result is in contrast to our own osteological height estimate observations in this study and those of multiple previous bioarchaeological studies^15,36,48^, which again may reflect potentially interesting inter-population variability as part of the nuanced complexity underlying subsistence shifts^92^.

One important potential limitation of our study, aside from the number of individuals, is uncertainty regarding the ultimate portability of polygenic scores over genetic, geographic, and temporal distances^81,82,95,96^. Our analyses were also based on incomplete ancient DNA genotype data (however, note that we did exclude potential error from deamination-based ancient DNA damage by masking all potentially affected sites in the primary versions of our analyses). Yet the significant, positive relationship between polygenic height scores and estimated osteological statures across our overall sample (Fig. 1F) demonstrates the biological plausibility of our model. Moreover, our primary results were unchanged when we incorporated archaeological site latitude and longitude variables into our analyses.

However, even with complete genome-wide genotype data, polygenic height scores only capture a proportion of the heritable component of stature variation^83,97–99^. Therefore, our primary analytical approach might incompletely capture the stature-relevant effects of any genetic ancestry variation across our sample. That is, with respect to our hypotheses, we would only be partially accounting for any cultural period-confounded migration/gene flow among populations with different genetic height profiles. For example, gene flow as a result of the spreading of the Yamnaya/Corded Ware cultures (“Steppe ancestries”) starting ~4,000 to 5,000 years ago may have been associated with the introduction of relatively greater proportions of ‘tall’ alleles into various regions of Europe^58,93^. If this is a general phenomenon that extends to small effect and other loci not included in our individual-level polygenic height score calculations, then our cultural period-related inferences could be erroneous.

Accordingly, to help explore the potential effects of these processes on our results we conducted a parallel set of analyses in which we included factors reflecting genome-wide genetic ancestry (from the first four axes of a multi-dimensional scaling analysis) in our model. To provide a sense of the relative magnitudes of potential impact of these effects, we observed that 21.3% of the variation in the difference between individual and per-sex mean osteological stature is explained by a model including both individual polygenic risk scores and the MDS components (SI Appendix; Table S14). In contrast, individual-level polygenic height score alone (i.e. without MDS factors) explained 4.35% of the variation (Fig. 1E).

When including the MDS factors in the analysis, our downstream finding of shorter osteological statures than expected (i.e., based on polygenic height scores and MDS factors, and in the context of the overall sample) in the Neolithic relative to other cultural periods was in fact partly attenuated. While this approach represents an over-correction (or, more precisely, some level of double accounting) via the inclusion of correlated genetic signals that cannot be deconvoluted readily due to the phenotype’s pervasive polygenicity, the findings nonetheless call for cautious interpretations of the main results in our study while awaiting expanded future sample sizes.

For a subset of the individuals in our study (n=91) we were additionally able to consider the extent to which three paleopathological markers of non-specific childhood and childhood-inclusive stress (linear enamel hypoplasia, cribra orbitalia, and porotic hyperostosis; each of which are maintained in the skeleton into adulthood) are associated with the relationship between osteological stature estimates and polygenic height scores. We observed a slight trend of relatively shorter than expected (given polygenic height score) adult statures among individuals with one or more childhood stress markers present (*SI Appendix*, Fig. S7). However, larger sample sizes will be necessary to more fully explore interplays among specific paleopathological indicators, osteological versus genetic height scores, and cultural periods. Still, our findings do at least suggest that factors underlying skeletal growth trajectories are separable, at least in part, from those leading to paleopathological indicators of stress. In particular, high rates of paleopathology are still observed in post-Neolithic cultural periods, for example in the Bronze Age, even after absolute osteological stature and actual versus expected stature averages have recovered.

In summary, we united previously disparate osteological and paleogenomic datasets for 160 prehistoric European individuals on a per-individual basis. Our results represent a novel advance in the study of whether and how a major cultural transition in human evolution affected physiological health. In particular, we show that the average Neolithic individual may have been relatively short even when correcting for expected individual genetic contributions to adult stature. This result may reflect reduced nutrition and/or increased infectious disease burden. We also preliminarily developed a framework for further consideration of these results in the context of particular paleopathological indicators of childhood stress. Looking forward, our model can be expanded in various dimensions, for example to different world regions or to more constrained spatial and temporal contexts, in order to further the study of emergent physiological trade-offs across periods of dramatic cultural or environmental change. Integrated osteological-genetic approaches will increasingly become important components of the toolkit for studying the dynamics of past human health.

## Materials and methods

### Processing ancient DNA sequence reads

Published sequence data were downloaded from Sequence Read Archive and European Nucleotide Archive databases as indicated per the respective papers alongside sequence data from in-process studies (*SI Appendix*, Table S1). FASTQ^65,67,1–103^ and BAM^66,68,69,104–117^ files from shotgun sequence (n=19) and DNA capture-based datasets (n=141) were aligned/realigned using the Burrows-Wheeler aligner (BWA) *aln*^118^ to the human reference genome (hg19, build 37) with seeding disabled. For unaligned shotgun sequence data (n=7)^65,100–103^, leeHom^119^ was used to trim adapters and merge reads. Reads from sequence libraries that were not treated to remove damage signatures typical of aDNA (“non-UDG”) were subject to rescaling using mapDamage 2.0^120^ after mapping^101,110^ which downscales quality values for likely aDNA misincorporations based on read position and damage pattern. For partial- and full-UDG-treated libraries, two base pairs at the 5’ and 3’ ends were trimmed (prior to mapping) using seqtk (https://github.com/lh3/seqtk) so as to not confound downstream analyses with potential post-mortem deamination at the terminal read ends^66,68,112,121^. SAMtools was used to sort mapped reads and filter for mapping quality 30 and minimum base pair (bp) 30, with duplicates removed using SAMtools *rmdup*^122^. Read groups were added using Picard’s AddOrReplaceReadGroups. Following the GATK workflow^123^, realigning indels was performed using RealignerTargetCreator and IndelRealigner, followed by BaseRecalibrator to minimize sequence error introduced by potential mismatches to the reference (github.com/smmarciniak/aDNA_osteo_height).

### Genotyping and imputation

We implemented GATK UnifiedGenotyper^123^ followed by imputation to maximize the amount of genetic information for downstream analyses and since genotype likelihood scores can be generated for imputation. We opted to impute diploid genotypes and missing sites for the individuals in our data set (using the 1000 Genomes Project Consortium^124^ reference panel) to minimize potential reference bias that may otherwise occur when using an alternative approach of randomly sampling one allele at each site^125^.

The 1000 Genomes Phase 3 genetic variants reference panel was used for genotyping and imputation, as provided by BEAGLE^126^ (ftp.1000genomes.ebi.ac.uk/vol1/ftp/release/20130502/). After removing variants that are not SNPs, multiallelic SNPs and X- and Y-chromosomes, 77,818,345 variants remained. UnifiedGenotyper^123^ was used to obtain genotypes and likelihood scores with the following parameters: --genotyping_mode --alleles <1000 Genomes reference panel>, --GENOTYPE_GIVEN_ALLELES, --output_mode EMIT_ALL_SITES, --AllSitesPLs -R <hg19 reference FASTA> (github.com/smmarciniak/aDNA_osteo_height). Due to the potential for post-mortem damage impacting C>T and G>A allele changes, the per chromosome VCF files of the called genotypes were filtered for potential post-mortem damage (modifying https://github.com/ryhui/imputation-pipeline/pmd_filter.py for multi-individual VCF files and removing deamination in both directions). Potential deamination signals C>T (T>C) and G>A (A>G) genotypes were replaced in ancient individuals heterozygous for these genotypes with ‘./.’ similar to previous paleogenomic studies^65,76^.

Genotype likelihoods were then estimated using the per-chromosome VCF files, followed by imputation of missing SNPs based on the genotype probability score using the 1000 Genomes phase 3 haplotypes (http://bochet.gcc.biostat.washington.edu/beagle/1000_Genomes_phase3_v5a/) and GRCh37 genomic maps (http://bochet.gcc.biostat.washington.edu/beagle/genetic_maps/). Parameters for estimating genotype likelihoods were: gprobs=true, gl=<input genotypes from UnifiedGenotyper> ref=<Beagle imputation reference panel>, map=<hg19 recombination map>. Imputation parameters were: gt=<GL_output_VCF>, gprobs=true, impute=true, ref=<Beagle imputation reference panel>, map=<GRCh37 recombination map> (github.com/smmarciniak/aDNA_osteo_height). This resulted in 30,761,499 markers imputed/genotyped across 160 individuals. Prior to downstream analyses, the imputed VCF was filtered for a minimum genotype probability of 0.99 to maximize confident genotype calls postimputation. We repeated the above pipeline of genotyping calling and imputation without filtering for potential deamination signals and the results were consistent with the deamination filtered data (*SI Appendix*, Fig. S1, Table S3).

### Assessing accuracy of imputation

We evaluated how sequence coverage may be impacting imputation accuracy (i.e., whether imputation is outperforming under high vs. low coverage conditions). We compared our imputed genotype data in the high coverage Loschbour individual (~16x)^104^ with downsampled BAM files (using SAMtools *-s* parameter^122^) from 5x coverage to 0.5x for chromosome 1 using SnpSift^127^. We obtained a concordance rate (in terms of total sites recovered) of approximately 97-99%, suggesting imputation accuracy is not dramatically lower in the low coverage imputed genotype data (*SI Appendix*, Table S16).

We also assessed the imputation accuracy of heterozygous sites by comparing the not imputed high coverage genotype data for Loschbour with each downsampled imputed BAM file from 0.1x to 3x coverage. At the lowest coverage of 0.3x, approximately 85% of heterozygous sites are recovered with subsequent increases of ~98% at 3x coverage (*SI Appendix*, Table S17).

### Polygenic height scores

Polygenic scores were computed by downloading a publicly available GWAS dataset from the UK Biobank^77^, specifically genome-wide summary statistics available from the Neale Lab (http://www.nealelab.is/uk-biobank, Round 2, accessed September 2018 and May 2019). The quality controls implemented for the publicly available UK Biobank dataset (e.g., MAF > 0.1%, HWE p-value > 1e-10 in 337,199 individuals) contains 10.8 million analyzable SNPs (http://www.nealelab.is/blog/2017/9/11/details-and-considerations-of-the-uk-biobank-gwas). The variant IDs (“variants.tsv.bgz”) were merged with the “standing height GWAS” file (“50_irnt.gwas.imputed_v3.both_sexes.tsv.bgz”), where the beta values represent the effect size of the “ALT” allele, subsequently used for performing the polygenic scores.

For our data, polygenic height scores were estimated using PLINK 1.9^128^ with clumping of independent SNPs. Clumping was used to identify the SNP with the lowest p-value in each LD block^128^. This approach retains SNPs with the strongest statistical evidence while reducing the correlation between the remaining SNPs^129^. Although all common SNPs could be used in polygenic scoring, clumping to remove SNPs that have limited statistical association is also a practical approach^129^. Clumping was performed at the genome-wide p-value 5e-08 using PLINK 1.9 parameters “--clump-r2 0.1”, “--clump-kb 1000” with the 1000 Genomes “Europeans” reference population panel to retain the most correlated SNPs (“index SNPs”) from the UK Biobank height summary statistics, which was then used to calculate the polygenic height scores. Polygenic scores were calculated using “--geno 0” to exclude missing genetic markers and the “-score” flag, extracting the specific index SNPs (github.com/smmarciniak/aDNA_osteo_height).

### Osteological data collection

The 160 individuals in our data set have a broad geographical, temporal, and cultural period ranges. Radiocarbon or archaeologically calibrated dates, latitude/longitude coordinates, genetic sex, and archaeological/cultural period were obtained from the original paleogenomic and archaeological publications of the paleogenomic data used for this study (*SI Appendix*, Table S1).

### Long bone measurement and stature reconstruction

Both newly-collected (n=86 individuals) and previously collected/published (n=54) osteological data were included in this study (*SI Appendix*, Table S1). For n=20 of the latter set of individuals, only pre-calculated terminal height estimates (based on unavailable long bone length measurements) were available (*SI Appendix*, Table S1). Only adult individuals were included in our study. For newly collected data this assessment was based on the complete fusion of all long bone epiphyses; we otherwise relied on classifications of ‘adult’ in the published record (which were likely based on the same criterion).

Permissions to collect new long bone measurement data were coordinated with researchers (coauthors on this publication) affiliated with the museums and university departments housing the various individuals. An osteometric board was used to measure the maximum length measurements (to the nearest millimeter) of the femur, tibia, radius, and humerus following standard osteological methods^44,130,131^. Intact long bones were selected, either the left or right side, depending on availability and preservation; if both sides were available and fully preserved, then both were measured.

We used a regression-based approach to reconstruct osteological stature from Ruff and colleagues^44^. These equations were developed using 501 individuals from across Europe ~7,000 BC-1900 AD, broadly approximating the geographical and temporal span of the individuals in our dataset. Sex-specific regression equations for the femur, tibia, humerus, and radius were used (*SI Appendix*, Table S2a), with standard error estimates ranging from 1.66% to 2.73%^44^. For the tibia, separate “north” and “south” equations are available^44^. Given the potential for migration occurring across the temporal and cultural periods in our data set we computed estimates from both equations and averaged them for the tibia-derived stature estimate for all individuals in our study (with available tibia measurements) regardless of geographic origin. When measurements from multiple different bones from the same individual were available, stature estimates derived from each of the different bones were estimated separately and then averaged to obtain a single point estimate per individual.

### Paleopathological indicators of non-specific stress

For paleopathological evaluations, 75 individuals were newly characterized and 16 were published/previously characterized (*SI Appendix*, Table S10). Crania with at least one permanent incisor were examined to record the presence or absence and severity of three skeletal indicators of non-specific stress: porotic hyperostosis, cribra orbitalia, and linear enamel hypoplasia^21,22,27,28^. Cribra orbitalia was assessed according to Stuart-Macadam (1991) on a five-stage scale of severity (n=74 evaluated) and whether the lesions were healed or active (n=73)^21^; porotic hyperostosis was evaluated on a three-stage scale (n=73) and whether the lesions were healed or active (n=72)^22^; and linear enamel hypoplasia was assessed as present or absent (i.e., whether one or more linear bands of decreased enamel thickness were visible) (n=76)^27^. The differences in the number of individuals assessed for healed or active lesions is the exclusion of a previously published individual (n=1 Neolithic)^29^ identified as having active lesions and n=1 Mesolithic individual for whom the nature of the porotic hyperostosis lesions was unspecified^132^.

### Statistical analyses

Statistical analyses were conducted in RStudio (v1.2.5033). Our main linear model was generated using osteological height and genetic height scores with sex as a co-variate. Data normality was assessed using ‘ggResidpanel’ (v0.3.0)^133^ (*SI Appendix*, Fig. S10). The residuals from this model were the basis for downstream analyses comparing patterns of stature variation across cultural periods as well as with the paleopathology data. Statistical analyses (t-tests) on the residuals from various linear models (including those below) were performed and the results are provided in the supplementary tables. False discovery rate calculations (p.adjust, method=“fdr”) were performed in R for each analysis set in order to help quantify the multiple testing effect.

We performed two additional analytical iterations of our linear model to evaluate consistency of downstream results. First, we included latitude and longitude as additional factors in the linear model framework described above. Second, we included factors related to genetic ancestry variation into our linear model, following the approach of Cox and colleagues^84^. Specifically, we performed a multi-dimensional scaling (MDS) analysis in PLINK (v1.9)^128^ using the full genomewide SNP genotype data available for all of the individuals in our study. We generated the “plink.genome” file (plink --file <input_plink_files> --genome) which was used as input for the MDS analysis specifying 4 dimensions (plink --file <input_plink_files> --read-genome plink.genome --cluster --mds-plot 4). The first four axes (C1, C2, C3 and C4) were each included in the linear model.

## Data availability

All osteological measurements, final stature estimates, and other skeletal individual-level information (e.g., ID, sex, radiocarbon dates, archaeological/ cultural period, geographical coordinates, publication sources for the ancient DNA data) are provided in Table S1 and additional supplementary tables. Although no new paleogenomic data were generated directly for this study, for n=28 individuals the analyzed ancient DNA data are from primary manuscripts currently in preparation or submitted for publication. References and accession numbers for these 28 individuals will be incorporated into updated versions of Table S1 as they are available. Scripts related to genotype calling, imputation, polygenic scoring, and statistical analyses are available at github.com/smmarciniak/aDNA_osteo_height.

## Acknowledgments

We thank many researchers and skeletal collections curators for their positive responses to us and their interest in this study even when materials or data were unavailable for inclusion. Components of this work were supported by the Harry J. and Elissa M. Sichi Early Career Professorship in Anthropology (to G.H.P.) and grants from the Wenner-Gren Foundation (#222377 to S.M.) and the National Institutes of Health (R01-GM115656 to G.H.P.). J.K. was supported by a long-term development project RV67985939 and by a grant of the Czech Science Foundation n. 19-20970Y. P.V. was supported by a grant from the Ministry of Culture of the Czech Republic (DKRVO 2019-2023/7.I.c, National Museum, 00023272). T.H., T.S.Z., K.K. were supported by a grant of the Hungarian Research, Development and Innovation Office (project number: FK128013 and K124326). G.H.P. thanks the DFG Center for Advanced Studies “Words, Bones, Genes, Tools” at the University of Tübingen for their support. Computations for this research were performed on the Pennsylvania State University’s Institute for Computational and Data Sciences’ supercomputing cluster. This content is solely the responsibility of the authors and does not necessarily represent the views of the Institute for Computational and Data Sciences. The osteological illustrations in Figures 1 and 2 were commissioned from Katharine Thompson.

## Supplementary Materials

**Figure S1.** Linear regressions and residuals of osteological height and genetic height score with sex as a co-variate without deamination filtering.

**Figure S2.** Residuals of osteological height and genetic height score with sex, latitude and longitude as co-variates.

**Figure S3.** Residuals of osteological height and genetic height score with sex as a co-variate, plotting females and males separately.

**Figure S4.** Replicability of the residuals of osteological height and genetic height score with sex as a co-variate using long bone lengths

**Figure S5.** Linear regressions of osteological height and genetic height score using long bone lengths.

**Figure S6.** Residuals of osteological height and genetic height score with sex and ancestries as co-variates.

**Figure S7.** Residuals of osteological height and genetic height score with sex as a co-variate for individuals with 1, 2, 3 paleopathological indicators of stress (n=53 individuals).

**Figure S8.** Residuals of osteological height and genetic height score with sex as a co-variate for individuals with healed cribra orbitalia and healed porotic hyperostosis.

**Figure S9.** Residuals of osteological height and genetic height score with sex as a co-variate for individuals with linear enamel hypoplasia and porotic hyperostosis across cultural periods (n=91 evaluated).

**Figure S10.** Diagnostic residual plots of the deamination filtered data.

**Table S1.** Description of individuals included in data set (n=160).

**Table S2a**. Average osteological heights across cultural periods.

**Table S2b.** Comparisons of the residuals from a linear model of osteological stature and sex.

**Table S3**. Comparisons of the residuals from a linear model of osteological stature and polygenic height score with sex as a co-variate for data not filtered for deamination.

**Table S4.** Polygenic height scores for deamination filtered and not deamination filtered data.

**Table S5.** Polygenic height score t-test results.

**Table S6.** Comparisons of the residuals from a linear model of osteological stature and polygenic height score with sex as a co-variate for deamination filtered data.

**Table S7.** Comparisons of the residuals from a linear model of osteological stature and polygenic height score with sex, latitude and longitude as co-variates.

**Table S8.** Comparisons of the residuals from a linear model of average long bone length and polygenic height score with sex as a co-variate for deamination-filtered data.

**Table S9.** Comparisons of residuals from a linear model of osteological stature and polygenic height score with sex and ancestries as covariates.

**Table S10.** Paleopathological summary for 91 individuals.

**Table S11**. Comparison of residuals from the main linear model for individuals with 0, 1+ and 2+ indicators of paleopathological stress (n=53 individuals).

**Table S12a.** Comparisons for individuals with LEH, cribra orbitalia or porotic hyperostosis with the residuals from a linear model of osteological and polygenic height score with sex as a covariate

**Table S12b.** Comparisons for individuals with LEH and healed cribra orbitalia or porotic hyperostosis with the residuals from a linear model of osteological height and polygenic height score with sex as a covariate.

**Table S13a.** Comparisons for individuals with LEH, cribra orbitalia or porotic hyperostosis with the residuals from a linear model of osteological and polygenic height score with sex as a covariate within cultural periods.

**Table S13b.** Comparisons for individuals with LEH and healed cribra orbitalia or porotic hyperostosis with the residuals from a linear model of osteological and polygenic height score with sex as a co-variate within cultural periods.

**Table S14a.** Comparisons for individuals with LEH, cribra orbitalia or porotic hyperostosis with the residuals from a linear model of osteological and polygenic height score with sex as a covariate across cultural periods.

**Table S14b.** Comparisons for individuals with LEH and healed cribra orbitalia or porotic hyperostosis with the residuals from a linear model of osteological and polygenic height score with sex as a co-variate across cultural periods.

**Table S15**. A model including both individual polygenic risk scores and MDS components

**Table S16.** Assessing imputation accuracy in high coverage vs. low coverage paleogenomic data

**Table S17.** Assessing imputation of heterozygote sites in imputed data.

## Notes

### Competing Interest Statement

The authors have declared no competing interest.

## References

1. Zeder, M. A. The origins of agriculture in the Near East. Curr. Anthropol. 52, 221 (2011).

2. Bar-Yosef, O. & Meadow, R. H. The origins of agriculture in the Near East. in Last hunters, first farmers: New perspectives on the prehistoric transition to agriculture (eds. Price, T. D. & Gebauer, A.-B.) 39–94 (School of American Research Press, 1995).

3. Pinhasi, R. & von Cramon-Taubadel, N. Craniometric data supports demic diffusion model for the spread of agriculture into Europe. PLoS One 4, e6747 (2009).

4. Bogucki, P. The spread of early farming in Europe. Am. Sci. 84, 242–253 (1996).

5. Fort, J. Demic and cultural diffusion propagated the Neolithic transition across different regions of Europe. J. R. Soc. Interface 12, (2015).

6. Gignoux, C. R., Henn, B. M. & Mountain, J. L. Rapid, global demographic expansions after the origins of agriculture. Proc. Natl. Acad. Sci. U. S. A. 108, 6044–6049 (2011).

7. Smith, B. D. The origins of agriculture in the Americas. Evol. Anthropol. Issues, News, Rev. 3, 174–184 (2005).

8. Page, A. E. et al. Reproductive trade-offs in extant hunter-gatherers suggest adaptive mechanism for the Neolithic expansion. Proc. Natl. Acad. Sci. 113, 4694–4699 (2016).

9. Lambert, P. M. Health versus fitness: competing themes in the origins and spread of agriculture? Curr. Anthropol. 50, 603–608 (2009).

10. Armelagos, G. J., Goodman, A. H. & Jacobs, K. H. The origins of agriculture: population growth during a period of declining health. Popul. Environ. 13, 9–22 (1991).

11. Zimmermann, A., Hilpert, J. & Wendt, K. P. Estimations of population density for selected periods between the Neolithic and AD 1800. Hum. Biol. 81, 357–380 (2009).

12. Bocquet-Appel, J.-P. The Neolithic Demographic Transition, Population pressure and Cultural change. Comp. Civilizations Rev. 58, 36–49 (2008).

13. Cohen, M. N. & Armelagos, G. J. Paleopathology at the Origins of Agriculture. (University Press of Florida, 1984).

14. Macintosh, A. A., Pinhasi, R. & Stock, J. T. Early life conditions and physiological stress following the transition to farming in Central/Southeast Europe: skeletal growth impairment and 6000 years of gradual recovery. PLoS One 11, e0148468 (2016).

15. Bennike, P. & Alexandersen, V. Population plasticity in southern Scandinavia: from oysters and fish to gruel and meat. in Health: Skeletal Indicators of Agricultural and Economic Intensification (eds. Cohen, M. N. & Crane-Kramer, G.) 130–148 (University Press of Florida, 2007).

16. Smith, P. & Horwitz, L. Ancestors and inheritors: a bioanthropological perspective on the transition to agropastoralism in the Southern Levant. in Ancient Health: Skeletal Indicators of Agricultural and Economic Intensification (eds. Cohen, M. N. & Crane-Kramer, G.) 207–222 (University Press of Florida, 2007).

17. Larsen, C. S. The agricultural revolution as environmental catastrophe: Implications for health and lifestyle in the Holocene. Quat. Int. 150, 12–20 (2006).

18. Larsen, C. S. Biological changes in human populations with agriculture. Annu. Rev. Anthropol. 24, 185–213 (1995).

19. The Backbone of History: Health and Nutrition in the Western Hemisphere. (Cambridge University Press, 2002).

20. Pinhasi, R. & Stock, J. T. Human bioarchaeology of the transition to agriculture. (John Wiley & Sons, 2011).

21. Stuart-Macadam, P. Anemia in Roman Britain: Poundbury Camp. in Health in Past Societies. Biocultural Interpretations of Human Skeletal Remains in Archaeological Contexts (eds. Bush, H. & Zvelebil, M.) 101–113 (BAR International Series No.567, 1991).

22. Stuart-Macadam, P. Porotic hyperostosis: Representative of a childhood condition. Am. J. Phys. Anthropol. 66, 391–398 (1985).

23. Rivera, F. & Mirazón Lahr, M. New evidence suggesting a dissociated etiology for cribra orbitalia and porotic hyperostosis. Am. J. Phys. Anthropol. 164, 76–96 (2017).

24. Walker, P. L., Bathurst, R. R., Richman, R., Gjerdrum, T. & Andrushko, V. A. The causes of porotic hyperostosis and cribra orbitalia: A reappraisal of the iron-deficiency-anemia hypothesis. Am. J. Phys. Anthropol. 139, 109–125 (2009).

25. Wapler, U., Crubézy, E. & Schultz, M. Is cribra orbitalia synonymous with anemia? Analysis and interpretation of cranial pathology in Sudan. Am. J. Phys. Anthropol. 123, 333–339 (2004).

26. Brickley, M. B. Cribra orbitalia and porotic hyperostosis: A biological approach to diagnosis. Am. J. Phys. Anthropol. 167, (2018).

27. Goodman, A. H., Armelagos, G. J. & Rose, J. C. Enamel hypoplasias as indicators of stress in three prehistoric populations from Illinois. Hum. Biol. 52, 515–528 (1980).

28. Goodman, A. H. & Rose, J. C. Dental enamel hypoplasias as indicators of nutritional status. in Advances in Dental Anthropology (eds. Kelley, M. & Larsen, C. S.) 279–293 (Wiley-Liss Inc., 1991).

29. Ash, A. et al. Regional differences in health, diet and weaning patterns amongst the first Neolithic farmers of central Europe. Sci. Rep. 6, 29458 (2016).

30. Holder, S., Miliauskienė, Ž., Jankauskas, R. & Dupras, T. An integrative approach to studying plasticity in growth disruption and outcomes: A bioarchaeological case study of Napoleonic soldiers. Am. J. Hum. Biol. e23457 (2020) doi:10.1002/ajhb.23457.

31. Dabbs, G. R. Health status among prehistoric Eskimos from Point Hope, Alaska. Am. J. Phys. Anthropol. 146, 94–103 (2011).

32. Eshed, V., Gopher, A., Pinhasi, R. & Hershkovitz, I. Paleopathology and the origin of agriculture in the Levant. Am. J. Phys. Anthropol. 143, 121–133 (2010).

33. Starling, A. P. & Stock, J. T. Dental indicators of health and stress in early Egyptian and Nubian agriculturalists: A difficult transition and gradual recovery. Am. J. Phys. Anthropol. 134, 520–528 (2007).

34. Zakrzewski, S. R. Variation in ancient Egyptian stature and body proportions. Am. J. Phys. Anthropol. 121, 219–229 (2003).

35. Formicola, V. & Giannecchini, M. Evolutionary trends of stature in Upper Paleolithic and Mesolithic Europe. J. Hum. Evol. 36, 319–333 (1999).

36. Holt, B. M. & Formicola, V. Hunters of the Ice Age: the biology of Upper Paleolithic people. Am. J. Phys. Anthropol. 137, 70–99 (2008).

37. Hoppa, R. D. & Saunders, S. R. Human Growth in the Past: Studies from Bones and Teeth. (Cambridge University Press, 1999).

38. Neves, W. & Wesolowski, V. Economy, nutrition, and disease in prehistoric coastal Brazil: a case study from the state of Santa Catarina. in The Backbone of History: Health and Nutrition in the Western Hemisphere (eds. Steckel, R. & Rose, J. C.) 376–400 (Cambridge University Press, 2002).

39. Vercellotti, G. et al. Exploring the multidimensionality of stature variation in the past through comparisons of archaeological and living populations. Am. J. Phys. Anthropol. 155, 229–242 (2014).

40. Eveleth, P. B. & Tanner, J. M. Worldwide variation in human growth. (Cambridge University Press, 1991).

41. Eveleth, P. B. Population differences in growth: environmental and genetic factors. in Human growth 373–394 (Springer, 1979).

42. Tanner, J. M. Introduction: Growth in height as a mirror of the standard of living. Stature Living Stand. Econ. Dev. Univ. Chicago Press. Chicago-London 1–6 (1994).

43. Steckel, R. H. Stature and the standard of living. J. Econ. Lit. 33, 1903–1940 (1995).

44. Ruff, C. B. et al. Stature and body mass estimation from skeletal remains in the European Holocene. Am. J. Phys. Anthropol. 148, 601–617 (2012).

45. Raxter, M. H., Ruff, C. B. & Auerbach, B. M. Technical note: revised Fully stature estimation technique. Am. J. Phys. Anthropol. 133, 817–818 (2007).

46. Vercellotti, G., Agnew, A. M., Justus, H. M. & Sciulli, P. W. Stature estimation in an early medieval (XI-XII c.) Polish population: Testing the accuracy of regression equations in a bioarcheological sample. Am. J. Phys. Anthropol. 140, 135–142 (2009).

47. Formicola, V. & Franceschi, M. Regression equations for estimating stature from long bones of early Holocene European samples. Am. J. Phys. Anthropol. 100, 83–88 (1996).

48. Sladek, V. et al. Central European Human Postcranial Variation. in Skeletal Variation and Adaptation in Europeans: Upper Paleolithic to the Twentieth Century (ed. Ruff, C. B.) 315–354 (Wiley-Blackwell, 2018).

49. Hermanussen, M. Stature of early Europeans. Hormones 2, 175–8 (2003).

50. Piontek, J. & Vančata, V. Transition to agriculture in Europe: Evolutionary trends in body size and body shape. Ecol. Asp. Past Hum. Settlements Eur. Bienn. Books EAA 2, 61–92 (2002).

51. Goodman, A. H., Lallo, J. W., Armelagos, G. J. & Rose, J. C. Health change at Dickson Mounds, Illinois (A.D. 950–1300). in Paleopathology at the Origins of Agriculture (eds. Cohen, M. N. & Armelagos, G. J.) 271–306 (Academic Press, 1984).

52. Walker, P. L. & Thornton, R. Health, nutrition, and demographic change in native California. in The Backbone of History: Health and Nutrition in the Western Hemisphereistory: health and nutrition in the Western Hemisphere (eds. Steckel, R. H. & Rose, J. C.) 506–523 (Cambridge University Press, 2002).

53. Lambert, P. M. Health in prehistoric populations of the Santa Barbara Channel Islands. Am. Antiq. 58, 509–522 (1993).

54. Temple, D. H. Patterns of systemic stress during the agricultural transition in prehistoric Japan. Am. J. Phys. Anthropol. 142, 112–124 (2009).

55. Pechenkina, E. A., Benfer, R. A. & Zhijun, W. Diet and health changes at the end of the Chinese Neolithic: The Yangshao/Longshan transition in Shaanxi province. Am. J. Phys. Anthropol. 117, 15–36 (2002).

56. Angel, J. L. Health as a crucial factor in the changes from hunting to developed farming in the eastern Mediterranean. in Paleopathology at the Origins of Agriculture (eds. Armelagos, G. J. & Cohen, M. N.) 51–74 (Academic Press, 1984).

57. Meiklejohn, C. & Key, P. Socioeconomic change and patterns of pathology and variation in the Mesolithic and Neolithic of Western Europe: Some suggestions. in Paleopathology at the Origins of Agriculture (eds. Cohen, M. N. & Armelagos, G. J.) 75–100 (University Press of Florida, 1984).

58. Cox, S. L., Ruff, C. B., Maier, R. M. & Mathieson, I. Genetic contributions to variation in human stature in prehistoric Europe. Proc. Natl. Acad. Sci. U. S. A. 116, 21484–21492 (2019).

59. Meiklejohn, C. & Zvelebil, M. Health status of European populations at the agricultural transition and the implications for the adoption of farming. Heal. past Soc. biocultural Interpret. Hum. Skelet. Remain. Archaeol. Context. 567, 129–144 (1991).

60. Lango Allen, H. et al. Hundreds of variants clustered in genomic loci and biological pathways affect human height. Nature 467, 832–838 (2010).

61. Silventoinen, K. Determinants of variation in adult body height. J. Biosoc. Sci. 35, 263–285 (2003).

62. Yang, J. et al. Common SNPs explain a large proportion of the heritability for human height. Nat. Genet. 42, 565–569 (2010).

63. Stulp, G. & Barrett, L. Evolutionary perspectives on human height variation. Biol. Rev. 91, 206–234 (2016).

64. Mathieson, I. et al. Genome-wide patterns of selection in 230 ancient Eurasians. Nature 528, 499–503 (2015).

65. Martiniano, R. et al. The population genomics of archaeological transition in west Iberia: Investigation of ancient substructure using imputation and haplotype-based methods. PLoS Genet. 13, e1006852 (2017).

66. Olalde, I. et al. The Beaker phenomenon and the genomic transformation of northwest Europe. Nature 555, 190–196 (2018).

67. Brace, S. et al. Ancient genomes indicate population replacement in Early Neolithic Britain. Nat. Ecol. Evol. 1 (2019) doi:10.1038/s41559-019-0871-9.

68. Mathieson, I. et al. The genomic history of southeastern Europe. Nature 555, 197–203 (2018).

69. Haak, W. et al. Massive migration from the steppe was a source for Indo-European languages in Europe. Nature 522, 207–211 (2015).

70. Larsen, C. S. et al. Bioarchaeology of Neolithic Çatalhöyük reveals fundamental transitions in health, mobility, and lifestyle in early farmers. Proc. Natl. Acad. Sci. 116, 12615–12623 (2019).

71. Holt, B. M. Mobility in Upper Paleolithic and Mesolithic Europe: Evidence from the lower limb. Am. J. Phys. Anthropol. 122, 200–215 (2003).

72. Larsen, C. S., Hutchinson, D. L. & Stojanowski, C. M. Health and lifestyle in Georgia and Florida: agricultural origins and intensification in regional perspective. in Ancient Health: Skeletal Indicators of Economic and Political Intensification (eds. Cohen, M. N. & Crane-Kramer, G. M. M.) 20–34 (University Press of Florida, 2007).

73. Cardoso, H. F. V. & Gomes, J. E. A. Trends in adult stature of peoples who inhabited the modern Portuguese territory from the Mesolithic to the late 20th century. Int. J. Osteoarchaeol. 19, 711–725 (2009).

74. Roberts, C. A. & Cox, M. The impact of economic intensification and social complexity on human health in Britain from 6000 BP (Neolithic) and the introduction of farming to the mid-nineteenth century AD. in Ancient Health: Skeletal Indicators of Agricultural and Economic Intensification (eds. Cohen, M. N. & Crane-Kramer, G. M. M.) 149–163 (University Press of Florida, 2007).

75. Hui, R., D’Atanasio, E., Cassidy, L. M., Scheib, C. L. & Kivisild, T. Evaluating genotype imputation pipeline for ultra-low coverage ancient genomes. Sci. Rep. 10, 18542 (2020).

76. Gelabert, P., Olalde, I., de-Dios, T., Civit, S. & Lalueza-Fox, C. Malaria was a weak selective force in ancient Europeans. Sci. Rep. 7, 1377 (2017).

77. Bycroft, C. et al. The UK Biobank resource with deep phenotyping and genomic data. Nature 562, 203–209 (2018).

78. Briggs, A. W. et al. Removal of deaminated cytosines and detection of in vivo methylation in ancient DNA. Nucleic Acids Res. 38, e87–e87 (2009).

79. Briggs, A. W. et al. Patterns of damage in genomic DNA sequences from a Neandertal. Proc. Natl. Acad. Sci. U. S. A. 104, 14616–14621 (2007).

80. Dabney, J., Meyer, M. & Pääbo, S. Ancient DNA damage. Cold Spring Harb. Perspect. Biol. 5, (2013).

81. Mostafavi, H. et al. Variable prediction accuracy of polygenic scores within an ancestry group. Elife 9, e48376 (2020).

82. Duncan, L. et al. Analysis of polygenic risk score usage and performance in diverse human populations. Nat. Commun. 10, 1–9 (2019).

83. Zaidi, A. A. & Mathieson, I. Demographic history mediates the effect of stratification on polygenic scores. Elife 9, 1–30 (2020).

84. Cox, S. L. et al. Predicting skeletal stature using ancient DNA. bioRxiv.

85. Mummert, A., Esche, E., Robinson, J. & Armelagos, G. J. Stature and robusticity during the agricultural transition: Evidence from the bioarchaeological record. Econ. Hum. Biol. 9, 284–301 (2011).

86. Larsen, C. S. Foraging to farming transition: global health impacts, trends, and variation. in Encyclopedia of Global Archaeology (ed. Smith, C.) 2818–2824 (Springer New York, 2014).

87. Milner, G. R. Early agriculture’s toll on human health. Proc. Natl. Acad. Sci. U. S. A. 116, 13721–13723 (2019).

88. Barker, G. & Richards, M. B. Foraging–farming transitions in Island Southeast Asia. J. Archaeol. Method Theory 20, 256–280 (2013).

89. Lander, F. & Russell, T. The archaeological evidence for the appearance of pastoralism and farming in southern Africa. PLoS One 13, e0198941 (2018).

90. Fuller, D. Q., Kingwell-Banham, E. J., Lucas, L., Murphy, C. & Stevens, C. Comparing pathways to agriculture. Archaeol. Int. 18, 61–66 (2015).

91. Cameron, M. E. The Riet River sites: Positioning regional diversity in the introduction of domesticated livestock to southern Africa. Journal of Archaeological Science: Reports vol. 23 72–79 (2019).

92. Rosenstock, E. et al. Human stature in the Near East and Europe ca. 10,000–1000 BC: its spatiotemporal development in a Bayesian errors-in-variables model. Archaeol. Anthropol. Sci. 11, 5657–5690 (2019).

93. Mathieson, I. et al. Genome-wide patterns of selection in 230 ancient Eurasians. Nature (2015) doi:10.1038/nature16152.

94. Niskanen, M., Ruff, C. B., Holt, B., Sladek, V. & Berner, M. Temporal and Geographic Variation in Body Size and Shape of Europeans from the Late Pleistocene to Recent Times. in Skeletal Variation and Adaptation in Europeans: Upper Paleolithic to the Twentieth Century 49–90 (2018).

95. De La Vega, F. M. & Bustamante, C. D. Polygenic risk scores: A biased prediction? Genome Med. 10, 100 (2018).

96. Martin, A. R. et al. Clinical use of current polygenic risk scores may exacerbate health disparities. Nat. Genet. 51, 584–591 (2019).

97. Sohail, M. et al. Polygenic adaptation on height is overestimated due to uncorrected stratification in genome-wide association studies. Elife 8, e39702 (2019).

98. Berg, J. J. et al. Reduced signal for polygenic adaptation of height in UK biobank. Elife 8, (2019).

99. Bitarello, B. D. & Mathieson, I. Polygenic Scores for Height in Admixed Populations. G3|Genes|Genomes|Genetics 10, 4027–4036 (2020).

100. González-Fortes, G. et al. Paleogenomic evidence for multi-generational mixing between Neolithic farmers and Mesolithic hunter-gatherers in the Lower Danube Basin. Curr. Biol. doi: 10.1016/j.cub.2017.05.023 (2017).

101. Cassidy, L. M. et al. Neolithic and Bronze Age migration to Ireland and establishment of the insular Atlantic genome. Proc. Natl. Acad. Sci. U. S. A. 113, 368–373 (2016).

102. Jones, E. R. et al. The Neolithic transition in the Baltic was not driven by admixture with early European farmers. Curr. Biol. 27, 576–582 (2017).

103. Skoglund, P. et al. Genomic diversity and admixture differs for Stone-Age Scandinavian foragers and farmers. Science. 344, 747–750 (2014).

104. Lazaridis, I. et al.Ancient human genomes suggest three ancestral populations for present-day Europeans. Nature 513, 409–413 (2014).

105. Olalde, I. et al.Derived immune and ancestral pigmentation alleles in a 7,000-year-old Mesolithic European. Nature 507, 225–228 (2014).

106. Sikora, M. et al. The population history of northeastern Siberia since the Pleistocene. Nature 570, 182–188 (2019).

107. Marcus, J. H. et al. Genetic history from the Middle Neolithic to present on the Mediterranean island of Sardinia. Nat. Commun. 11, 1–14 (2020).

108. Furtwängler, A. et al. Ancient genomes reveal social and genetic structure of Late Neolithic Switzerland. Nat. Commun. 11, 1–11 (2020).

109. Rivollat, M. et al. Ancient genome-wide DNA from France highlights the complexity of interactions between Mesolithic hunter-gatherers and Neolithic farmers. Sci. Adv. 6, eaaz5344 (2020).

110. Jones, E. R. et al. Upper Palaeolithic genomes reveal deep roots of modern Eurasians. Nat. Commun. 6, doi:10.1038/ncomms9912 (2015).

111. Lipson, M. et al. Parallel palaeogenomic transects reveal complex genetic history of early European farmers. Nature 551, 368–372 (2017).

112. Mittnik, A. et al. The genetic prehistory of the Baltic Sea region. Nat. Commun. 9, 442 (2018).

113. Allentoft, M. E. et al. Population genomics of Bronze Age Eurasia. Nature 522, 167–172 (2015).

114. Keller, A. et al. New insights into the Tyrolean Iceman’s origin and phenotype as inferred by whole-genome sequencing. Nat. Commun. 3, doi: 10.1038/ncomms1701 (2012).

115. Seguin-Orlando, A. et al. Genomic structure in Europeans dating back at least 36,200 years. Science. 346, 1113–1118 (2014).

116. Sikora, M. et al. Ancient genomes show social and reproductive behavior of early Upper Paleolithic foragers. Science. 358, 659–662 (2017).

117. Fu, Q. et al. The genetic history of Ice Age Europe. Nature 534, 200–205 (2016).

118. Li, H. & Durbin, R. Fast and accurate short read alignment with Burrows-Wheeler transform. Bioinformatics 25, 1754–1760 (2009).

119. Renaud, G., Stenzel, U. & Kelso, J. leeHom: adaptor trimming and merging for Illumina sequencing reads. Nucleic Acids Res. 42, e141–e141 (2014).

120. Jónsson, H., Ginolhac, A., Schubert, M., Johnson, P. L. F. & Orlando, L. mapDamage2.0: fast approximate Bayesian estimates of ancient DNA damage parameters. Bioinformatics 29, 1682–1684 (2013).

121. Gamba, C. et al. Genome flux and stasis in a five millennium transect of European prehistory. Nat. Commun. 5, 5257 (2014).

122. Li, H. et al. The Sequence Alignment/Map format and SAMtools. Bioinformatics 25, 2078–2079 (2009).

123. McKenna, A. et al. The Genome Analysis Toolkit: a MapReduce framework for analyzing next-generation DNA sequencing data. Genome Res. 20, 1297–303 (2010).

124. Auton, A. et al. A global reference for human genetic variation. Nature vol. 526 68–74 (2015).

125. Günther, T. & Nettelblad, C. The presence and impact of reference bias on population genomic studies of prehistoric human populations. PLOS Genet. 15, e1008302 (2019).

126. Browning, S. R. & Browning, B. L. Rapid and accurate haplotype phasing and missingdata inference for whole-genome association studies by use of localized haplotype clustering. Am. J. Hum. Genet. 81, 1084–97 (2007).

127. Cingolani, P. et al. A program for annotating and predicting the effects of single nucleotide polymorphisms, SnpEff: SNPs in the genome of Drosophila melanogaster strain w1118; iso-2; iso-3. Fly (Austin). 6, 80–92 (2012).

128. Purcell, S. et al. PLINK: A Tool Set for Whole-Genome Association and Population-Based Linkage Analyses. Am. J. Hum. Genet. 81, 559–575 (2007).

129. Marees, A. T. et al. A tutorial on conducting genome-wide association studies: Quality control and statistical analysis. Int. J. Methods Psychiatr. Res. 27, e1608 (2018).

130. Ruff, C. Skeletal Variation and Adaptation in Europeans: Upper Paleolithic to the Twentieth Century. (Wiley-Blackwell, 2018).

131. Buikstra, J. E. & Ubelaker, D. H. Standards for Data Collection from Human Skeletal Remains: Proceedings of a Seminar at the Field Museum of Natural History. (Arkansas Archeological Survey, 1994).

132. Serrulla Rech, F. & Sanin Matias, M. Forensic anthropological report of Elba. (2017).

133. Goode, K. & Rey, K. ggResidpanel (v0.3.0). (2019).

