## Supplementary Figures_S1-S10 for "An integrative skeletal and paleogenomic analysis of prehistoric stature variation suggests relatively reduced health for early European farmers"

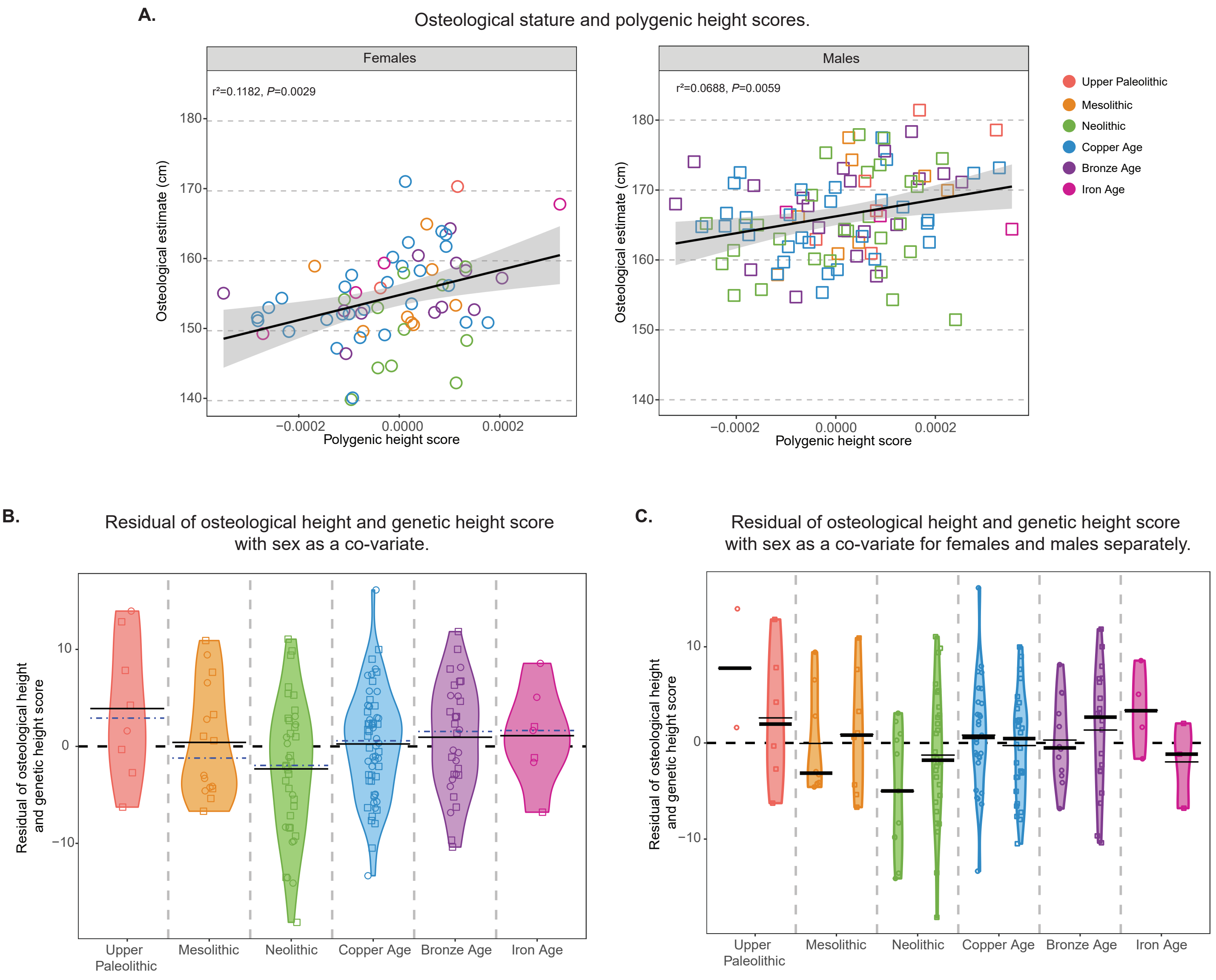


**Fig S1.** Linear regressions and residuals of osteological height and genetic height score with sex as a co-variate without deamination filtering. **A)** The relationship between polygenic height score and estimated osteological stature (cm) for females and males. **B)** Residuals of the relationship between polygenic height score and osteological height with sex as a co-variate for all individuals, by cultural period. Mean and median are represented by the black and blue dashed lines, respectively. **C)** Residuals of the relationship between polygenic height score and osteological height with sex as a co-variate for females and males plotted separately. Mean is represented by a thin line and median by the rectangle. Females are represented by circles and males by squares. Full results in Table S3.

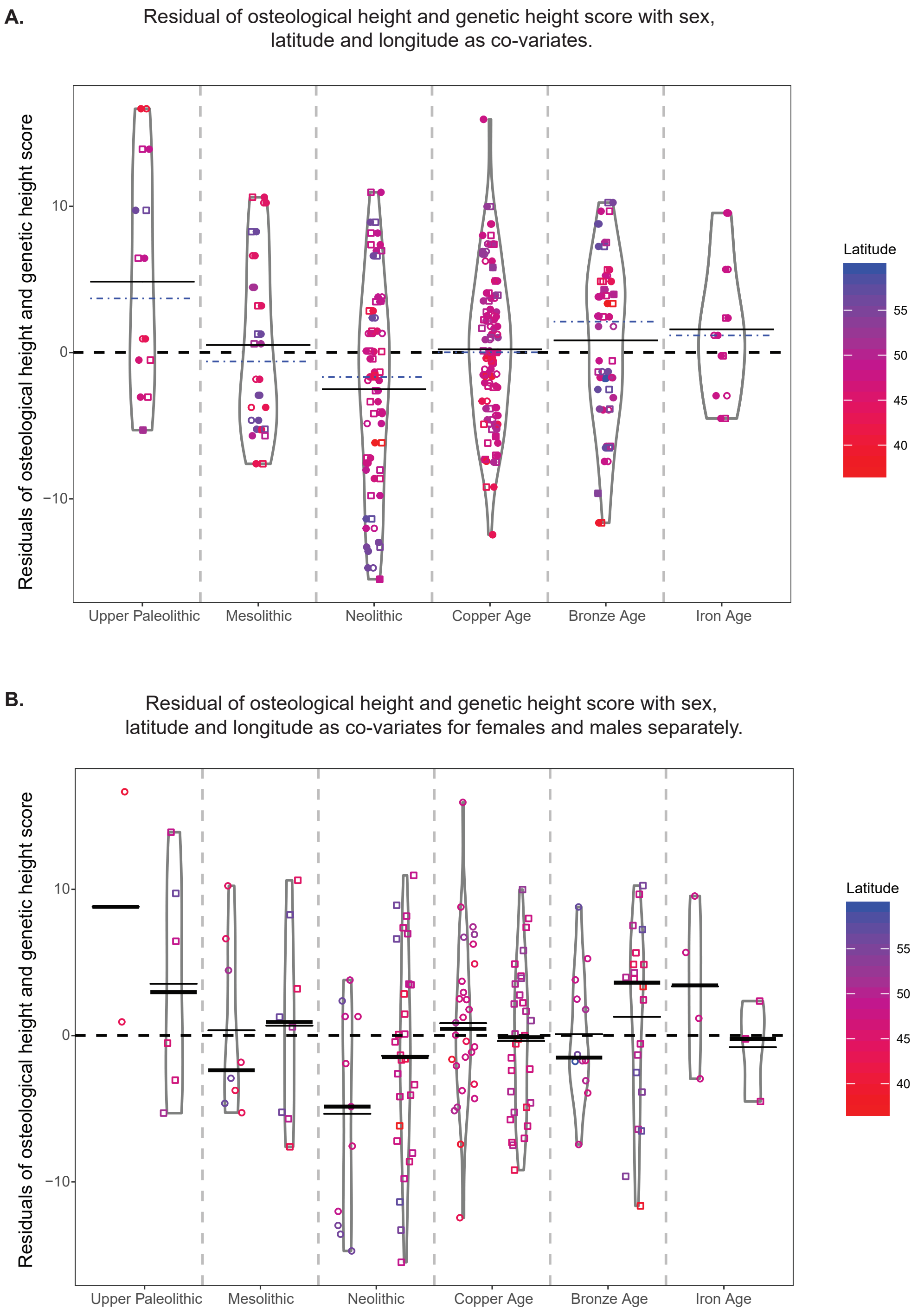


**Fig. S2.** Residuals of osteological height and genetic height score with sex, latitude and longitude as co-variates. Latitude and longitude were included as co-variates in the main linear model to assess replicability of the main results. **A**) For females and males combined, Pre-Neolithic individuals (average residual = +1.96 ± 6.9 cm) were ~4.47 cm taller than expected relative to Neolithic individuals (average residual = -2.50 ± 7.2 cm; P=0.018; FDR=0.062). The average osteological vs. genetic height score residual then increased steadily in the Copper Age (+2.72 cm relative to the Neolithic; P=0.047; FDR=0.063), Bronze Age (+3.34 cm; P=0.031; FDR=0.062), and Iron Age (+4.08 cm; P=0.085; FDR=0.085). **B**) Females and males represented separately across cultural periods. Latitude gradient (north to south) is indicated. Mean is represented by a thin line and median by the rectangle. Females are represented by circles and males by squares. Full results in Table S7.

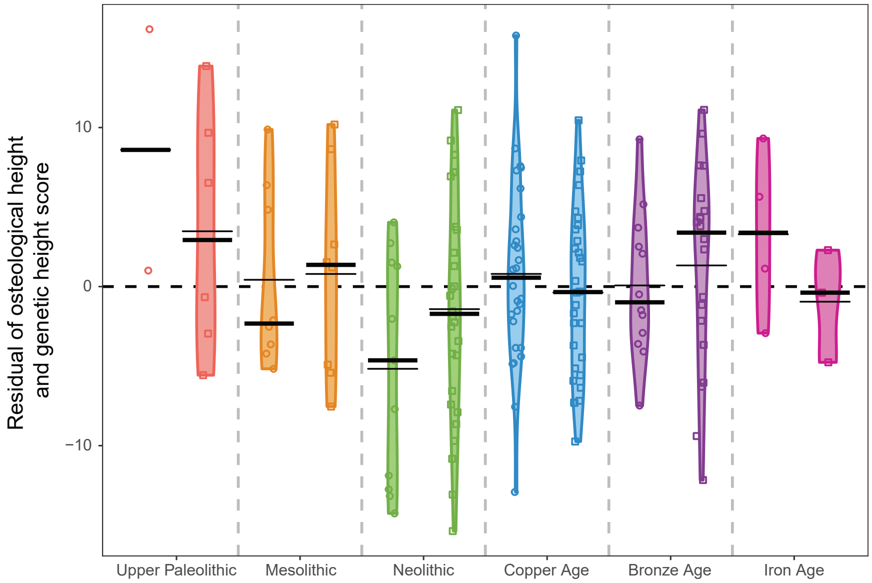


**Fig. S3.** Residuals of osteological height and genetic height score with sex as a co-variate. Females (circles) and males (squares) are plotted side by side based on the residuals of the relationship between polygenic height score and osteological height with sex as a co-variate. Mean is represented by the thin line and median by the rectangle. Full results in Table S6.

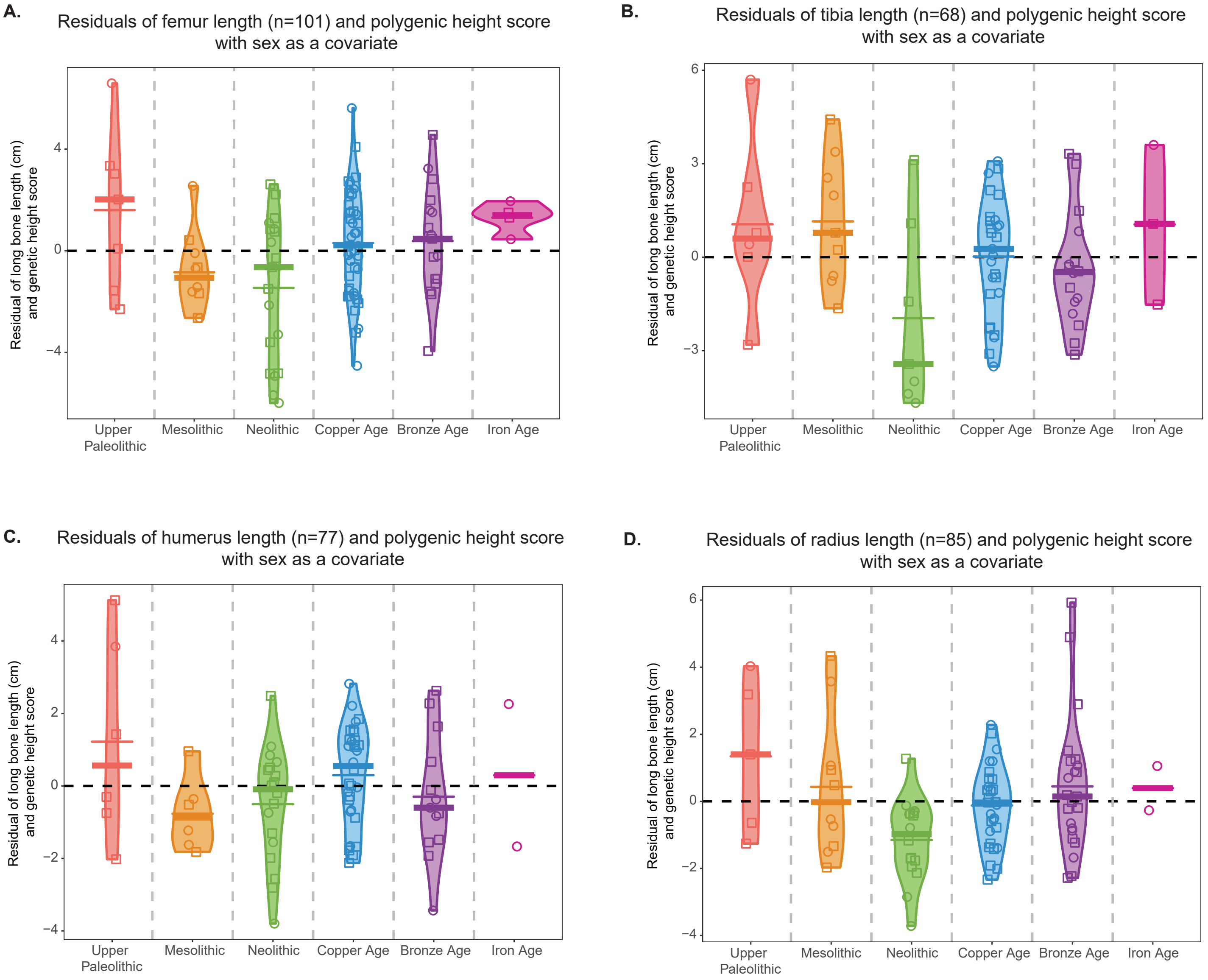


**Fig. S4.** Replicability of the residuals of osteological height and genetic height score with sex as a co-variate using long bone lengths. Residuals of the relationship between polygenic height score and long bone lengths for the femur (**A**), tibia (**B**), humerus (**C**) and radius (**D**) with sex as a co-variate for all individuals, by cultural period. Mean is represented by the thin line and median by the rectangle. Females are represented by circles and males by squares. Full results in Table S8.

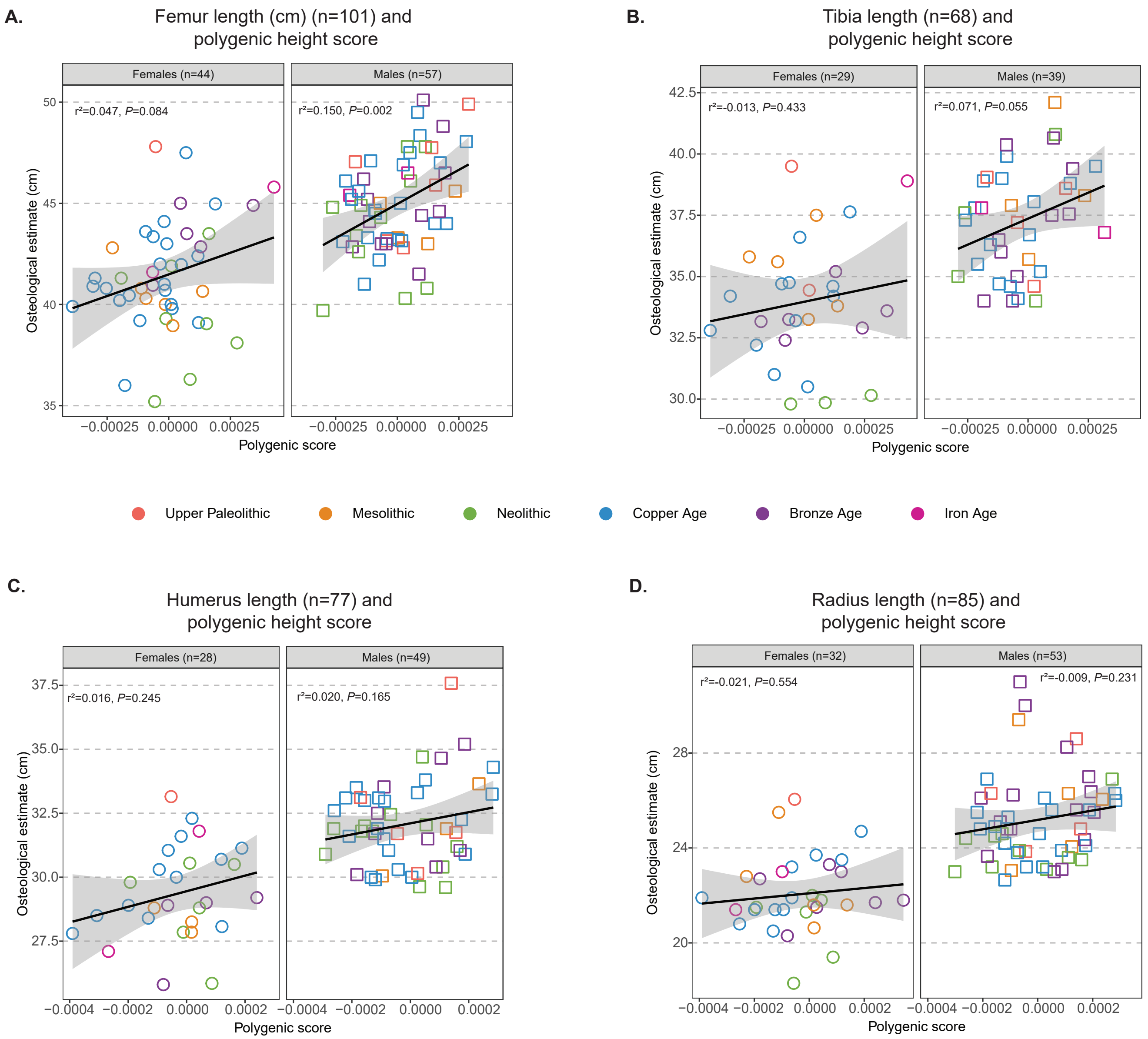


**Fig. S5.** Linear regressions of osteological height and genetic height score using long bone lengths. The relationship between polygenic height score and long bone length (cm) for females and males for the femur (**A**), tibia (**B**), humerus (**C**) and radius (**D**) by cultural period. Females are represented by circles and males by squares.

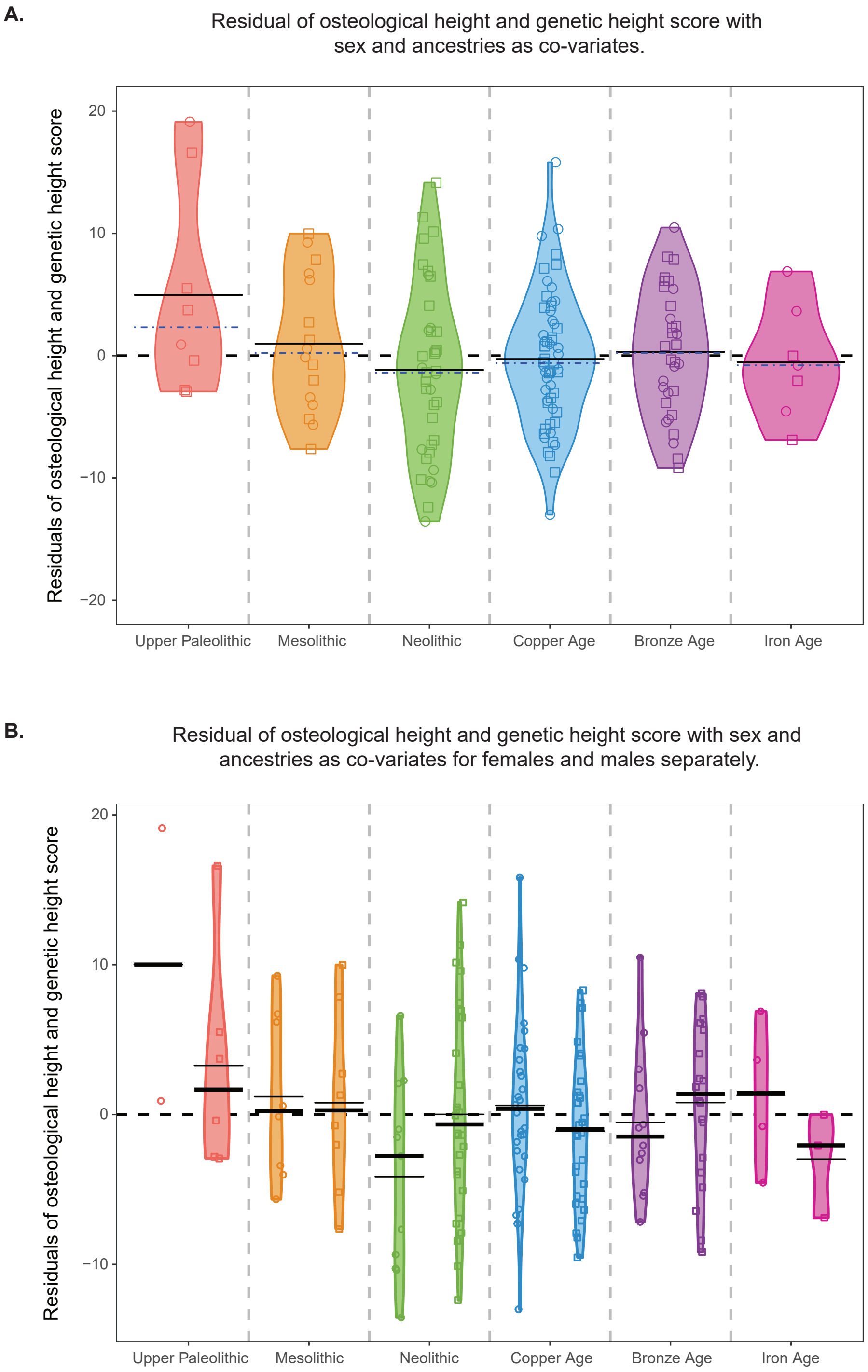


**Fig. S6**. Residuals of osteological height and genetic height score with sex and ancestries as co-variates. Genetic ancestries based on the MDS clusters are included as co-variates in the main linear model. **A**) Females and males combined, mean is in black and median in blue dashed line. **B**) Females and males represented separately, with mean as the thin line and median as the rectangle. Females are represented by circles and males by squares. Full results in Table S9.

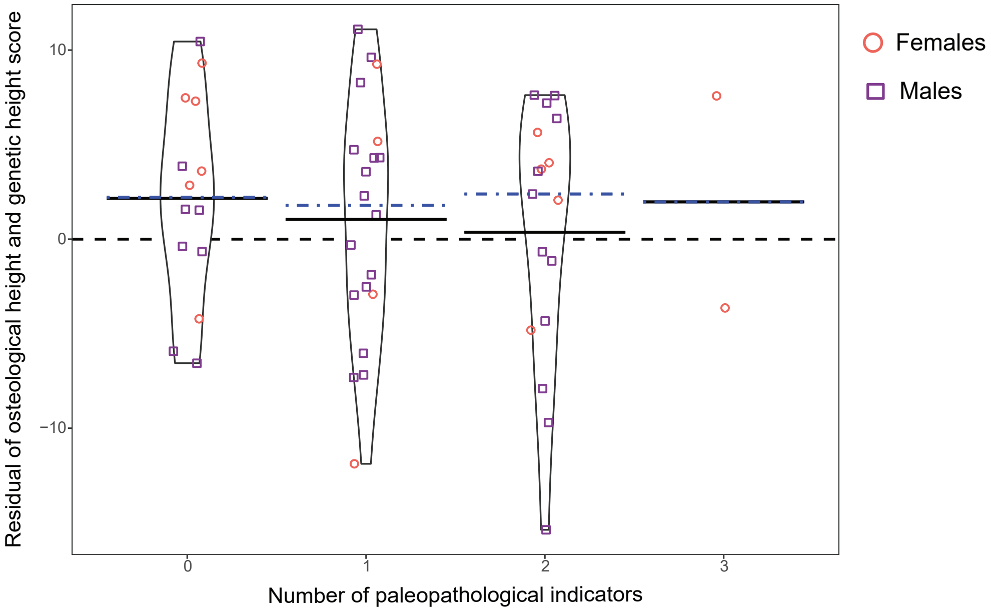


**Fig. S7.** Residuals of osteological height and genetic height score with sex as a co-variate for individuals with 1, 2, 3 paleopathological indicators of stress (n=53 individuals). Residuals of osteological height and genetic height score with sex as a co-variate for 53 individuals who could be assessed for all three non-specific stress indicators. Individuals with 1, 2, 3 paleopathological indicators of stress are represented for females (circles) and males (squares). The mean is represented by the black line and the median the blue dashed line.

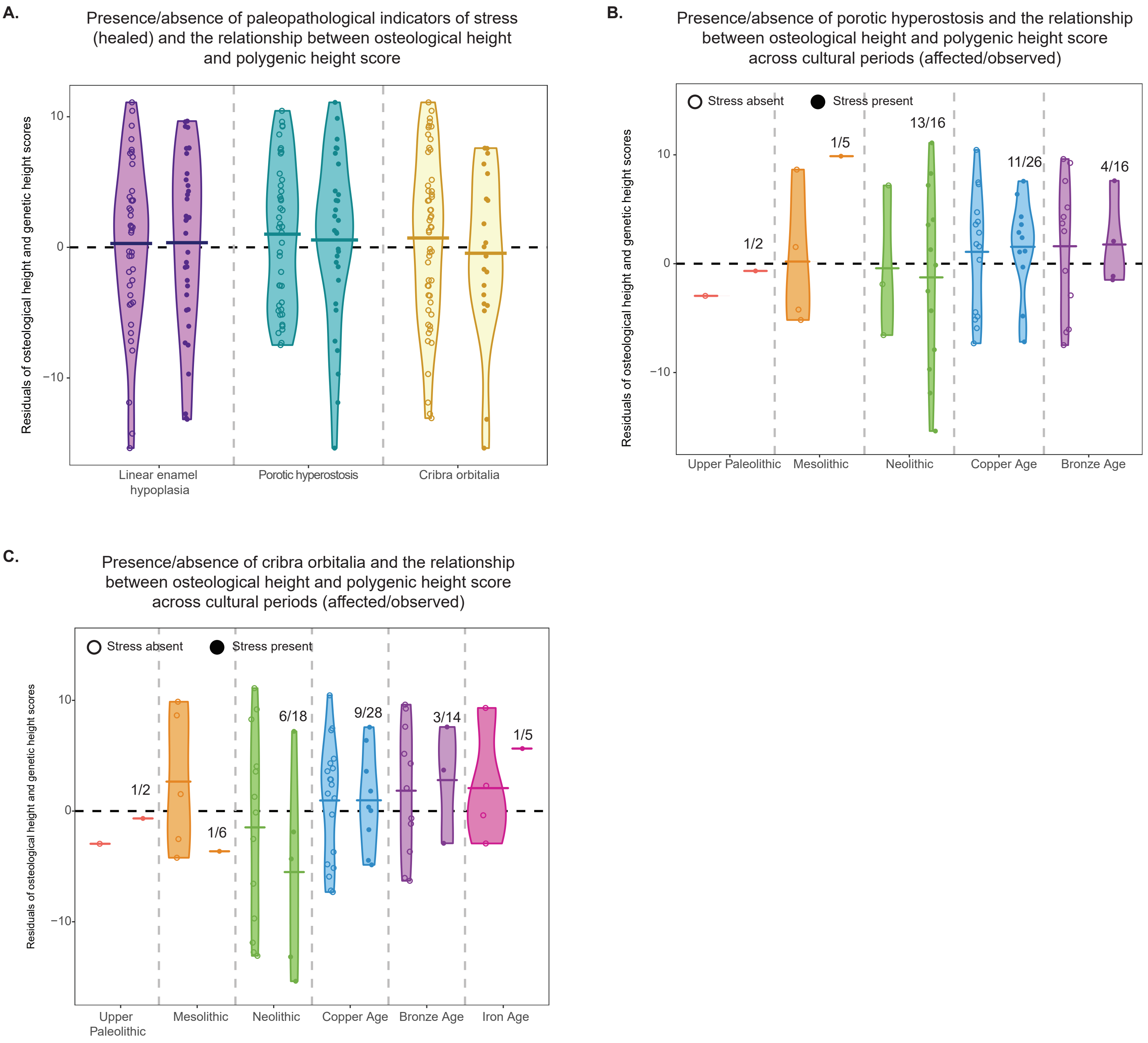


**Fig. S8.** Residuals of osteological height and genetic height score with sex as a co-variate for individuals with healed cribra orbitalia and healed porotic hyperostosis. **A**) Residuals of osteological height and genetic height score with sex as a co-variate are compared to individuals with healed cribra orbitalia and healed porotic hyperostosis. The data is also represented across cultural periods for porotic hyperostosis (**B**) and cribra orbitalia (**C**). Means are represented by the thin lines. Numbers above the bars indicate number of individuals. Full results in Table S12.

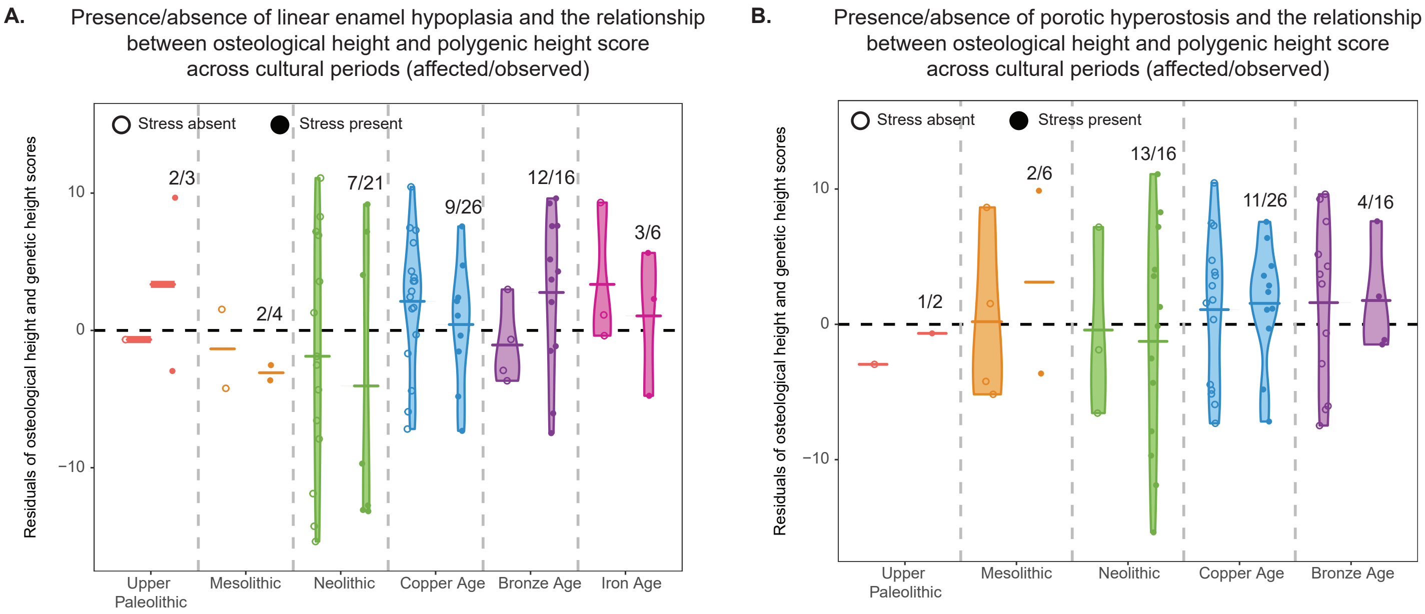


**Fig. S9.** Residuals of osteological height and genetic height score with sex as a co-variate for individuals with linear enamel hypoplasia and porotic hyperostosis across cultural periods. Comparison of the residuals generated from the main linear model to individuals with paleopathological indicators across cultural periods. Means are represented by the thin lines. Numbers above the bars indicate number of individuals. Full results in Table S13.

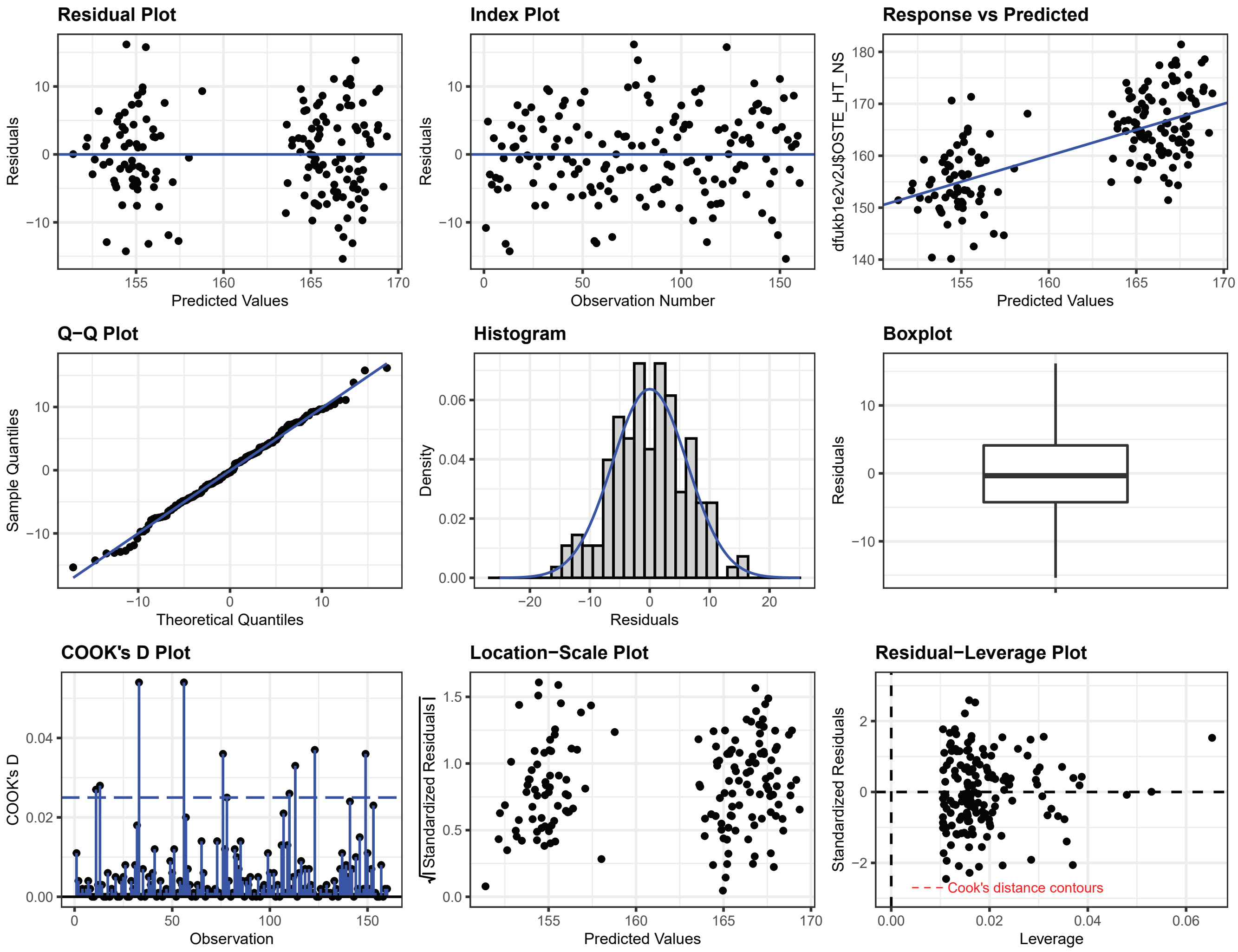


**Fig. S10.** Diagnostic residual plots for the linear model of osteological height and genetic height score with sex as a co-variate. Residual diagnostic plots generated using ‘ggResidpanel’ (v0.3.0) for the deamination filtered data set.
